# Natural genetic variation drives microbiome selection in the *Caenorhabditis elegans* gut

**DOI:** 10.1101/2021.03.12.435148

**Authors:** Fan Zhang, Jessica L. Weckhorst, Adrien Assié, Ciara Hosea, Christopher A. Ayoub, Anastasia Khodakova, Mario Loeza Cabrera, Daniela Vidal, Marie-Anne Félix, Buck S. Samuel

**Affiliations:** Alkek Center for Metagenomics and Microbiome Research and Department of Molecular Virology and Microbiology, Baylor College of Medicine, Houston TX; Program in Quantitative and Computational Biosciences, Baylor College of Medicine, Houston TX; Program in Development, Disease Models and Therapeutics, Baylor College of Medicine, Houston TX; Ecole Normale Supérieure, IBENS, CNRS UMR8197, INSERM U1024, Paris, France

**Keywords:** host-microbe interactions, genetics, gnotobiotic models, insulin signaling, model microbiome

## Abstract

Host genetic landscapes can shape microbiome assembly in the animal gut by contributing to the establishment of distinct physiological environments. However, the genetic determinants contributing to the stability and variation of these microbiome types remain largely undefined. Here, we use the free-living nematode *Caenorhabditis elegans* to identify natural genetic variation among wild strains of *C. elegans* strains that drives assembly of distinct microbiomes. To achieve this, we first established a diverse model microbiome that represents the phylogenetic and functional diversity naturally found in the *C. elegans* microbiome. Using this community, we show that *C. elegans* utilizes immune, xenobiotic and metabolic signaling pathways to favor the assembly of different microbiome types. Variations in these pathways were associated with the enrichment for specific commensals, including the Alphaproteobacteria *Ochrobactrum*. Using RNAi and mutant strains, we showed that host selection for *Ochrobactrum* is mediated specifically by host insulin signaling pathways. *Ochrobactrum* recruitment is blunted in the absence of *daf-2*/IGFR and requires the insulin signaling transcription factors *daf-16/*FOXO and *pqm-1*/SALL2. Further, the ability of *C. elegans* to enrich for *Ochrobactrum* is correlated positively with host outcomes, as animals that develop faster are larger and have higher gut *Ochrobactrum* colonization as adults. These results highlight a new role for the highly conserved insulin signaling pathways in the regulation of microbiome composition in *C. elegans*.

## INTRODUCTION

Across kingdoms, shifts in microbiome composition accompany and contribute to host development, health, and physiology ^1–3^. Along with diet and lifestyle, host genetics can regulate the size and composition of the microbiome ^4–7^. This is apparent in human diseases with altered microbiome composition such as inflammatory bowel disease and obesity ^6^. While predicted host polymorphic loci for the development of these diseases have been identified ^8^, the majority of the data linking gut microbiome composition and host disease remains correlative. Thus, there is a great need to identify causal genetic host determinants that contribute to the stability and variation of microbiome types in order to effectively develop microbiome interventions as potential therapies.

To address this problem, we used wild strains of the nematode *Caenorhabditis elegans* and established a new diverse 63-member model microbiome that better represents the phylogenetic and functional diversity of the *C. elegans* wild microbiome, termed ‘BIGbiome’. This system proves several key advantages. *C. elegans* itself has a transparent body plan and robust genetic toolbox, and its short lifespan and amenability to high-throughput methods increase experimental throughput ^9,10^. *C. elegans* also shares many conserved pathways with higher organisms that could regulate microbiome recruitment, including metabolic, stress and innate immune pathways ^11–13^. Yet, it has been difficult to determine which of these pathways may contribute to microbiome community outcomes because much of our current understanding comes from decades of studies of the *C. elegans* lab strain N2-Bristol in association with *Escherichia coli* or human pathogens ^14^. By contrast, wild *C. elegans* strains encounter a large variety of microbes and selectively recruit only some of these from the environment to form its gut microbiome ^15–17^. This process is akin to the horizontal transmission of microbes that occurs throughout the animal kingdom, but one that remains poorly understood overall. Thus, probing host genetics with a representative natural microbial community may more completely reveal causal host determinants that contribute to microbiome outcomes.

We make progress towards this goal here. We present the most comprehensive examination to date of the causal influence of *C. elegans* natural genetic variation on the establishment of its gut microbiome. To achieve this, we utilized our newly developed model microbiome to test natural variation in its acquisition by a panel of nearly 40 ‘germ-free’ wild strains of *C. elegans* ^18^. We found that these strains selected for and acquired one of only three distinct types of gut microbiomes: (i) one dominated by *Ochrobactrum*, (ii) another dominated by *Bacteroidetes*, and (iii) one similar in composition to the bacterial lawn. Selection of these microbes was robust and consistent within the host strain, suggestive of a deterministic process driven by host genetic variation. To probe this variation, we conducted phenotypic, genetic and transcriptional profiling of wild strains representative of each microbiome type. This analysis revealed fundamental differences in host immune, stress and metabolic responses specific for the acquisition of each microbiome. Genetic gain and loss function studies further reveal a key role for insulin signaling pathways in microbiome regulation. In particular, we identify a previously uncharacterized role for insulin signaling in wild strains of *C. elegans* in the promotion of selective acquisition and maintenance of the gut microbiome via a DAF-2/PQM-1 pathway. Finally, we find that the ability of particular host genetic backgrounds to acquire a given microbiome directly influences host health fate. Higher levels of insulin signaling and broad activation of immune pathways promoted intestinal acquisition of otherwise rare *Ochrobactrum* from the bacterial lawns, and this was associated with faster growth rates. In contrast, low levels of insulin signaling activated non-selective stress responses and resulted in gut microbiomes that resemble the lawn and were associated with lower rates of host growth. Together, these studies establish an approach that utilizes wild worms and their natural microbes for microbiome studies and uses this system to identify novel roles for host insulin signaling in positive selection of pro-growth microbial communities.

## RESULTS

### Establishment of a diverse, representative core microbiome of *C. elegans*

Effective identification of host genes that drive assembly of distinct microbiomes requires a diverse model microbial community that closely resembles the variation a host may encounter in the wild. We reasoned that such a community should be: (i) reflective of the major microbial taxa found in natural microbiomes of wild *C. elegans*; (ii) highly functionally redundant; and (iii) easy to use and create in the lab. To achieve this, we expanded on previous analyses of the core microbiome of wild *C. elegans* populations ^19^ and selected bacterial strains from our collections (>500 strains) that matched the 14 core families of the *C. elegans* microbiome— *Enterobacteriaceae, Pseudomonadaceae, Xanthomonadaceae, Sphingomonadaceae, Sphingobacteriaceae, Flavobacteriaceae, Weeksellaceae, Acetobacteraceae, Moraxellaceae, Oxalobacteraceae, Comamonadaceae, Rhodobacteraceae, Microbacteriaceae,* and *Actinomycetales*. The resulting community, termed BIGbiome001 (referred to as ‘BIGbiome’ hereafter), comprises 63 strains from 23 genera (10 of 14 core families). Together, it represents 50-80% of the biomass in natural microbiomes of wild *C. elegans* [**Table 1**; see strain origins in **Table S1**; **Figure S1A-B**]. We also sought to model the functional redundancy observed in natural communities by including several taxonomically related bacterial strains from distinct wild *C. elegans* strains or habitats (range of 2-22 strains). BIGbiome complements simplified microbiomes like the recently developed 12-member CeMbio community ^20^ as it reflects the extensive strain level microbial diversity found in *C. elegans* natural microbiomes while still remaining experimentally tractable.

**Table 1.**
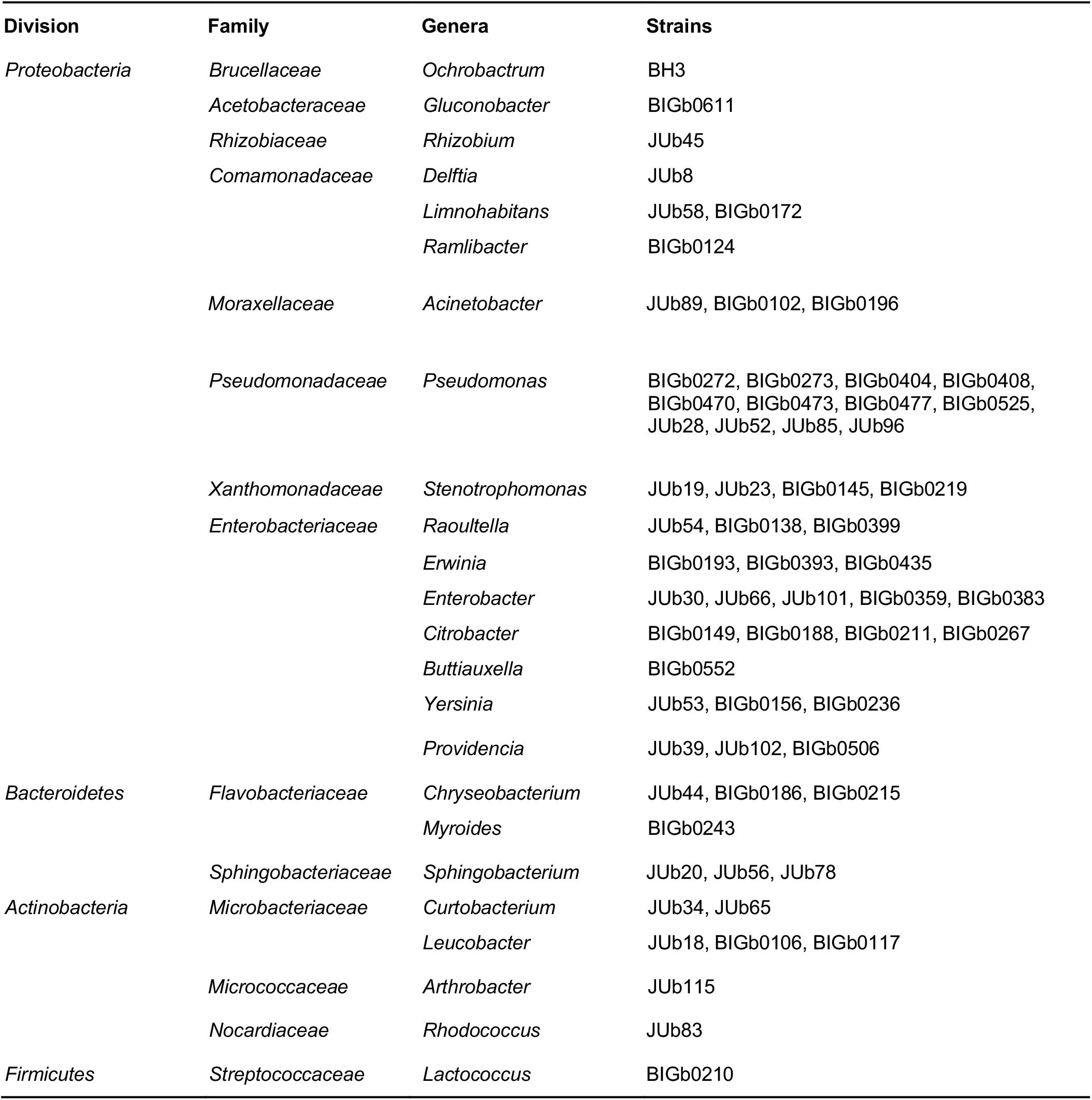
Summary of microbial strains in BIGbiome model microbiome.

### Development of distinct gut microbiome types in adult *C. elegans*

We next tested the robustness of the BIGbiome community for microbiome studies by profiling how it is acquired in the lab strain (N2) and a wild *C. elegans* strain (JU1218). To achieve this, we established a phenotyping pipeline for microbiome-based measures that include gut colonization density and composition [**Figure 1A**]. These methods allow for high-throughput determination of both the levels of overall bacterial colonization and proportions of bacteria that colonize the *C. elegans* gut. In this approach, strains are first made ‘germ-free’ by bleaching eggs followed by synchronization at the L1 stage. L1 animals are then exposed to the BIGbiome community (proportional mixture of each strain) on agar plates and monitored over their development and into adulthood. Using this approach, we found that appreciable *C. elegans* gut colonization could not be observed until day 1 of adulthood (48 hours post-L1). Differences between N2 and JU1218 host strains were observed by day 3 of adulthood and appeared to stabilize at that time [**Figure S1C-D**]. These results are consistent with previous studies of bacterial colonization of the *C. elegans* N2 intestine by *E. coli* ^21^. For these reasons, we chose days 1 and 3 of adulthood for further studies.

**Figure 1.**
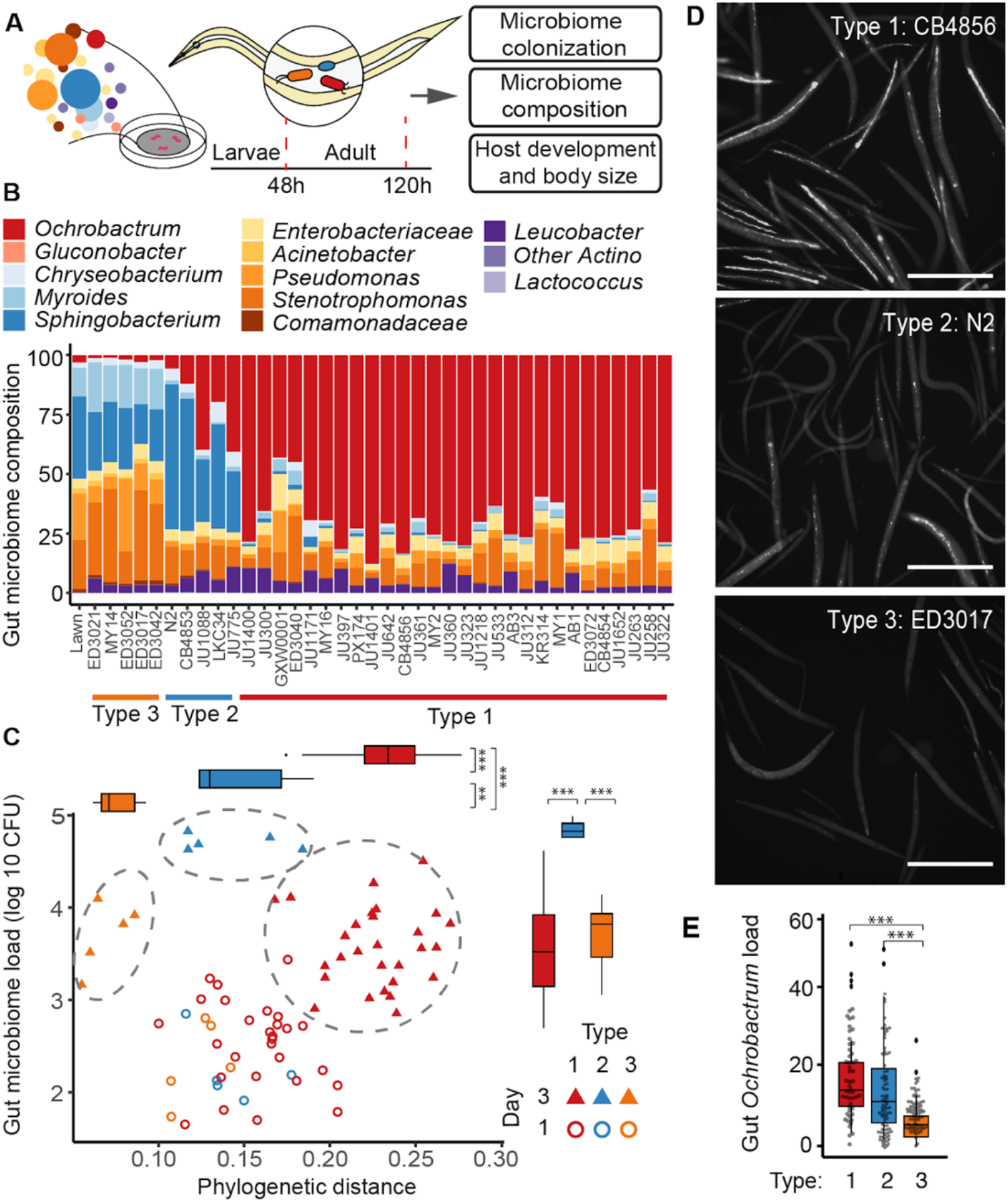
Natural genetic variation in *C. elegans* drives distinct gut microbiome types. **A**. Schematic diagram illustrating the pipeline to measure gut microbiome and host phenotypes of 38 *C. elegans* strains grown on microbiome mixture. Worm samples were collected at 48 h (day 1 adults) and 120 h (day 3 adults) after exposing synchronized L1 populations to BIGbiome. **B**. Gut microbiome composition of the 38 *C. elegans* strains in day 3 adulthood. Relative microbiome abundance was presented here as the mean of biological duplicates for each strain. **C**. The 38 strains were clustered into three distinct microbiome types based on their gut microbiome load per animal (y-axis) and phylogenetic distances to BIGbiome lawn (x-axis). Solid symbols showed samples collected in day 3 adulthood and open symbols showed samples in day 1 adulthood. Inset: Box-whisker plot of microbiome load per animal in three microbiome types. Type 2 strains (n=10) carried significantly higher gut microbiome load than Type 1 (n=56) and Type 3 strains (n=10). Box-whisker plot of phylogenetic distances to BIGbiome for the three microbiome types. Phylogenetic distances between each strain and BIGbiome lawn were calculated by weighted UniFrac. Type 1 strains (n=56) showed further distance to the BIGbiome lawn than Type 2 (n=10) and Type 3 strains (n=10). n represents the number of independent worm populations. **D**. Representative images of *C. elegans* strains in day 3 adulthood from each of the three microbiome types grown on BIGbiome with an isogenic GFP expressing *Ochrobactrum* strain. Bar = 500 μm. **E**. Box-whisker plot of GFP intensity quantified from fluorescent images of *C. elegans* strains grown on BIGbiome (GFP-*Ochrobactrum*) showed higher GFP-*Ochrobactrum* colonization in Type 1 (n=55) and 2 (n=94) strains than Type 3 strains (n= 73). n, individual animals quantified by microscopic images; P-values were generated from one-way ANOVA, followed by Tukey posthoc test with 95% confidence level and adjusted for multiple comparisons (***p<0.001, **p<0.01).

To assess the impact of host genetic variation on microbiome selection, we next used a genetically tractable but diverse host community. We selected 38 well characterized, fully genome sequenced and genetically distinct *C. elegans* strains ^18^ [**Table S2**]. Together with the lab strain (N2), these wild strains were first made ‘germ-free’ by bleaching eggs, and synchronized L1 animals were exposed to the BIGbiome community on agar plates. Animals from each strain were collected in bulk as adults at early (day 1) and later (day 3) stages of microbiome establishment and then assayed for differences in microbiome composition and gut colonization density. All of the worm strains exhibited both low levels of colonization and a comparable, lawn-like composition of their gut microbiomes at day 1 [**Figure S2**]. By day 3 of adulthood, however, the gut microbiomes became largely distinct from the surrounding bacterial lawn, and hosts exhibited up to a 30-fold range in levels of colonization [**Figure 1, Figure S2**].

Through our analysis of these microbiome communities, we found that the majority of the strains within the BIGbiome community colonized the guts of at least two independent worm strains [91.6%, 55 strains]. However, the assemblages and proportions of microbes observed in each worm strain were unique. This is similar to other systems ^22,23^ and may reflect the functional redundancy of the isolates within the microbiome. For example, *Enterobacteriaceae* exhibit significant genomic plasticity and are common in wild *C. elegans* microbiomes ^19,24^. Though resolution of this family in our samples is limited due to high identity of small subunit (SSU) rRNA genes, *Enterobacteriaceae* were consistent colonizers as a group (5-10% relative abundance). Other consistent colonizers included *Pseudomonas*, *Stenotrophomonas*, and *Comamonas* [**Table S3**]; the more rare *Leucobacter* was also enriched 30-60 fold in the worm gut relative to the bacterial lawn [**Table S3**]. Thus, within this community we observe taxa that are general colonizers and those that exhibit more strain-specific (see below) or stochastic colonization of the *C. elegans* gut [**Figure S3A-B**].

We next asked whether particular microbiome representations were favored more than others. We performed weighted UniFrac-based clustering of the animals by microbiome types on day 3 of adulthood and found that gut microbiome composition robustly separated into three microbiome types. Clustering was driven by dominant microbial taxa, and we termed the host clusters as Type 1, 2, or 3 [**Figure 1A-B, Figure S4A-B**]. The Type 1 hosts contained the largest group of *C. elegans* strains, harboring 28 strains. Notably, these strains were dominated by *Ochrobactrum pituitosum BH3* [>40% relative abundance; **Figure 1B**], a microbe previously identified as a common beneficial member of the *C. elegans* microbiome in the wild ^25–27^. The microbiomes of Type 2 strains (lab strain N2 and four wild strains) were dominated by several Bacteroidetes taxa (e.g., *Myroides*, *Chryseobacterium* and *Sphingobacterium*). These animals displayed reduced levels of *Ochrobactrum* in the gut [10-40% relative abundance; **Figure 1B**] and higher levels of gut colonization overall [2 to 30-fold higher than Type 1 or Type 3 strains; **Figure 1C, Figure S2C**]. Type 3 animals (five wild worm strains) were nearly devoid of gut *Ochrobactrum* and instead were dominated by high levels of *Bacteroidetes, Pseudomonas* and *Stenotrophomonas* [**Figure 1B**]. Overall, the microbiome of Type 3 strains resembled that of the bacterial lawn [**Figure 1C, Figure S4**]. Machine learning-based (random forest) classification of the most discriminant taxa supports this categorization scheme [Accuracy = 84.9%] and identified *Ochrobactrum*, *Sphingobacterium* and several rarer members as robust classifiers of the microbiome types [**Figure S3C**]. Analyses of overall gut microbiome alpha-diversity within samples indicates that the dominance of *Ochrobactrum* in the Type 1 strains had a tempering impact on microbiome diversity [Faith’s phylogenetic diversity of 6.3±0.9 in Type 1 vs. 8.9±0.7 in Type 3; Figure S4C]. Enrichment of otherwise rare microbes like *Ochrobactrum* from the lawn in the gut microbiome also increased beta-diversity between samples. The highest enrichment differential between the host microbiome composition relative to the lawn was observed for Type 1 animals, with more moderate enrichment observed in Types 2 and 3 [**Figure 1C, Figure S4D**].

We next utilized the inherent transparency of *C. elegans* to assess microbial enrichment on a single animal basis in order to examine individual variation within a given host strain. To accomplish this, we created BIGbiome mixtures where *Ochrobactrum BH3* was replaced by an isogenic GFP-expressing strain. Both microscopy- and large particle flow cytometry (Biosorter)-based analyses supported our finding that *Ochrobactrum* enrichment was greater in individuals from Type 1 strains (CB4856), particularly when compared to Type 3 animals (ED3017) that have limited *Ochrobactrum* colonization [P<0.0001; **Figure 1DE**]. Type 2 animals (N2 or LKC34) exhibited a broader distribution of *Ochrobactrum* levels on a per animal basis [**Figure 1E**]. This may be due to an inherent stochasticity in microbial levels and composition during the colonization process, as has been shown for the lab strain of *C. elegans* (N2) under certain conditions ^28^. Together, our results highlight three robust modes of microbiome regulation by host strains that vary in their selectivity for the microbes that colonize and their relative levels within the gut.

### *C. elegans* growth rates and body size correlate with adult microbiome composition

Previous studies have shown that exposure to individual microbes can have dramatically different impacts on the physiology and development of *C. elegans* ^13,16^. For example, many Alphaproteobacteria and *Enterobacteriaceae* strains generally promote growth, while most *Bacteroidetes* and *Stenotophomonas* strains are detrimental to N2 worms ^16,17^. Many of these same bacterial strains are present within the BIGbiome community. We thus examined representative *C. elegans* strains from each microbiome type for changes in growth rates or body sizes after development. Each of the strains were grown on agar plates containing either BIGbiome or *E. coli* OP50 lawns from L1 until adulthood (46-58hrs). Notably, all strains tested exhibited faster growth rates on the BIGbiome community compared to those grown on *E. coli* OP50 alone [**Figure 2A-B**]. The extent of the growth promotion did differ by microbiome type, however. Type 1 strains (JU1400) exhibited 75% faster growth versus 65% and 40% for Type 2 and 3 strains, respectively [**Figure 2A-B**]. Type 3 (ED3017) animals were also significantly smaller than the other microbiome types after 48hrs of development, although these differences did normalize by day 3 of adulthood [**Figure 2C, Figure S5A**]. Both faster developmental growth rates and/or larger body sizes at the L4 stage correlated with higher gut colonization of *Ochrobactrum* [Pearson of 0.74 and 0.49, respectively, P<0.002; **Figure 2D-E**] and lower *Enterobacter* [∼3% relative abundance; Pearson of 0.45 with body size only, P<0.005] and *Leucobacter* colonization [∼5% relative abundance; Pearson = 0.49 with growth rate only, P<0.005; Table S4]. Conversely, slower growth rates and smaller body size were associated with more permissive colonization by nine other genera: Bacteroidetes (*Chryseobacterium* and *Myroides*), Betaproteobacteria (*Limnohabitans, Ramlibacter,* and *Delftia*), Gammaproteobacteria (*Acinetobacter* and *Stenotrophomonas*), and Actinobacteria (*Arthrobacter* and *Curtobacterium*) [P<0.05; Table S4]. No significant correlations were observed between the overall gut microbiome load and either growth rates or body size [**Figure S5B-C**]. These studies indicate that host-driven acquisition of a selected microbial community during development can both positively and negatively alter host outcomes in adulthood.

**Figure 2.**
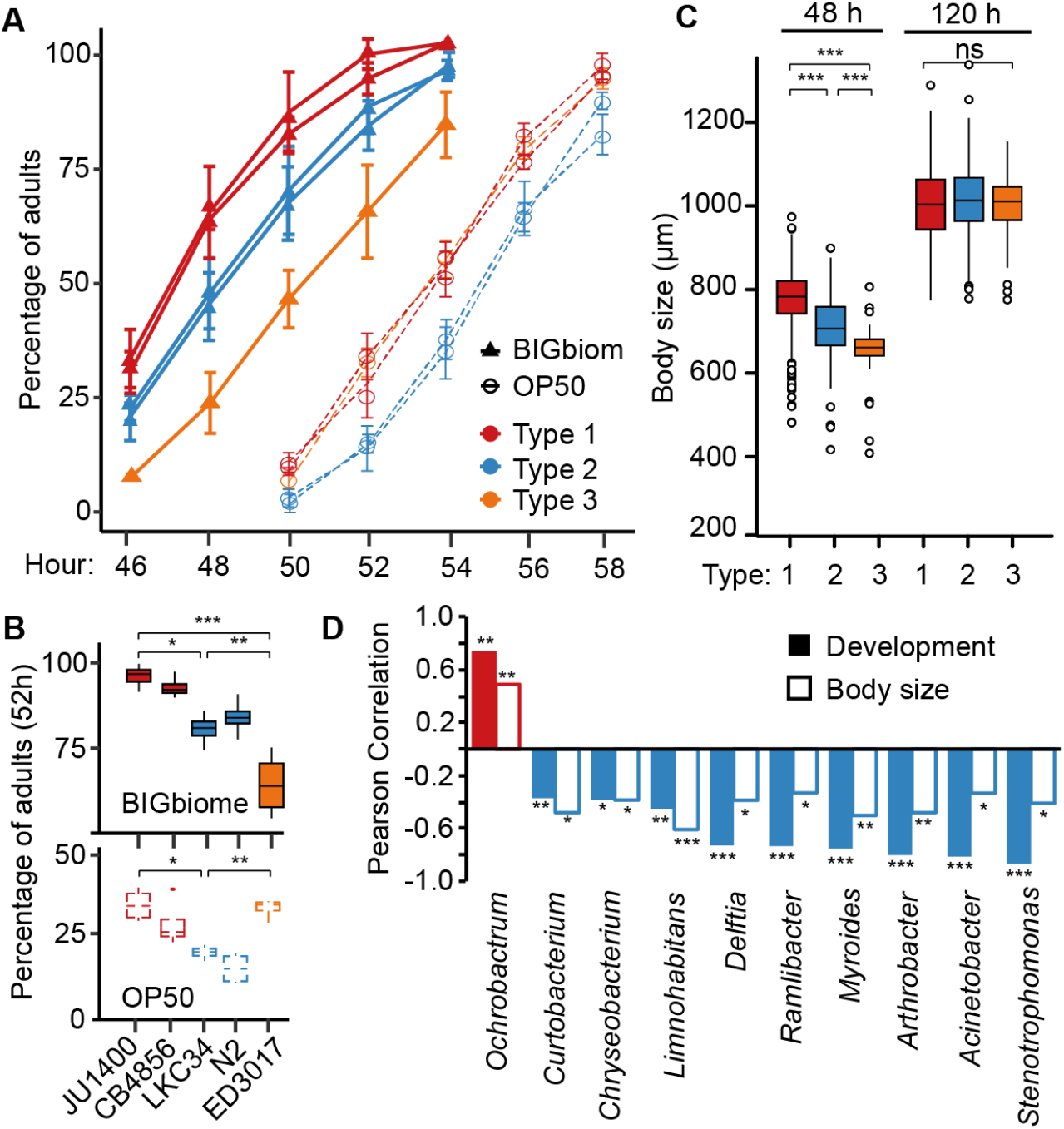
*C. elegans* developmental growth rates and body size during development correlate with adult microbiome. **A.** Developmental growth rates of representative strains from each microbiome type [Type 1 (JU1400 and CB4856), Type 2 (N2 and LKC34) and Type 3 (ED3017)] grown on BIGbiome and *E. coli* OP50. Percentage of adults are represented as mean ± SD with 4 replicates for each condition; representative of 3 independent experiments. **B.** Box-whisker plots of percent adults at 52 h post L1 stage (from A). Number of individual animals: BIGbiome (JU1400: n=145, CB4856: n=141, N2: n=132, LKC34: n=176, ED3017: n=157); *E. coli* OP50 (JU1400: n=173, CB4856: n=166, N2: n=142, LKC34: n=142, ED3017: n=153). **C.** Box-whisker plot of *C. elegans* body size by microbiome types at 48 h and 120 h post L1 stage. Type 1 strains (n=1076) had longer body size than Type 2 (n=168) and Type 3 strains (n=158) at 48 h. No significant difference among Type 1 (n=501), 2 (n=68), and 3 (n=56) at 120 h. P-values (for B and C) were generated from: one-way ANOVA, followed by and post hoc Tukey Honest Significant Difference test with 95% confidence level and adjusted for multiple comparisons (*** p<0.001, ** p<0.01,* p<0.05, n.s not significant). **D.** Pearson correlations of microbial taxa abundance (day 3 adults) with host developmental rates (52 h post L1) and body size (48 h post L1). The test statistic is based on Pearson’s product moment correlation coefficient and follows a t distribution with length(x)-2 degrees of freedom at the level of 95% confidence interval. *Ochrobactrum* (colored in red) is the only microbial taxa with positive correlation with both host phenotypes (p<0.05). 9 microbial taxa (colored in blue) show negative correlations with both host phenotypes (p<0.05).

### Natural genetic variation is associated with microbiome composition within the *C. elegans* gut

Our results indicate that the potential for specific natural variation in host genetics may drive the selection of particular microbiome communities. To explore these host pathways, we used the extensive genomics resources available through the Million Mutation Project for our wild *C. elegans* panel [>3.8M single nucleotide variants, ∼65,000 missense mutations versus N2 reference genome ^18^. We performed GWAS analyses (see *Methods*) to identify regions of the genome associated with taxa abundance, colonization level, alpha diversity and beta diversity as trait values per strain. We identified several regions that were associated with taxa abundance of *Chryseobacterium*, *Enterobacteriaceae*, *Gluconobacter*, *Acinetobacter*, *Curtobacterium* and *Leucobacter* across host strains [17-56% variance explained per taxa (relative and/or absolute abundance); 1308 total genes in 9 loci; **Figure S6**; see full list in **Table S5**]. As a whole, nine loci were enriched for genes with previously unknown functions [419 genes; Q=0.0039; WormCat tool ^29^], suggesting that microbiome studies may help ascribe phenotypes for these genes in the future. Notably, the most significant overlap was observed for genes that are upregulated in insulin receptor (*daf-2*/IGFR) mutants [316 genes; Q=1.3e-32 to dataset ^30^; WormExp tool ^31^]. Together, these analyses indicate that natural genetic variation may drive microbiome compositional differences.

### Type 1 animals express a broad repertoire of microbial response pathways to create selectivity

To more specifically identify the host signaling networks regulating selection of the gut microbiome, we transcriptionally profiled the host responses to colonization of a panel of representative *C. elegans* strains from each of the three microbiome types [Type 1, JU1400 and ED3040; Type 2, N2, LKC34, and CB4853; and Type 3, ED3017, MY14, and ED3042] [**Figure 3A, Table S6**]. At day 3 of adulthood, we observed a large set of differentially expressed genes between Types 1 and 3 [1507 higher in Type 1 (‘Type 1 Up’), 1706 higher in Type 3 (‘Type 3 Up’); **Figure 3B, Table S6**], consistent with the differences in microbiome composition between these strains. We first tested for correlations between transcript and taxa abundance across all of the *C. elegans* strains. We identified 2844 genes that were differentially expressed by microbiome type and significantly correlated with taxa abundance of one or more microbes [**Figure 3C**]. Interestingly, *Ochrobactrum-*correlated genes dominated the taxa-specific signatures, and genes that were positively correlated with *Ochrobactrum* were negatively correlated with Bacteroidetes *Myroides* (259 genes) and vice versa (857 genes). Smaller subsets of genes were correlated with the abundance of 16 other taxa, and these genes sets were largely distinct from those of *Ochrobactrum* and *Myroides* [**Figure 3C**]. These data could indicate that similar transcriptional networks coordinate the enrichment *Ochrobactrum* and the exclusion of *Myroides*. To begin to identify the function of these and other host genes that were upregulated in association with particular microbial communities, we used the WormExp tool ^31^. We observed broad increases in expression in genes involved in three main pathways: microbial and immune response, general stress response, and insulin signaling [**Figure 3D-G**].

**Figure 3.**
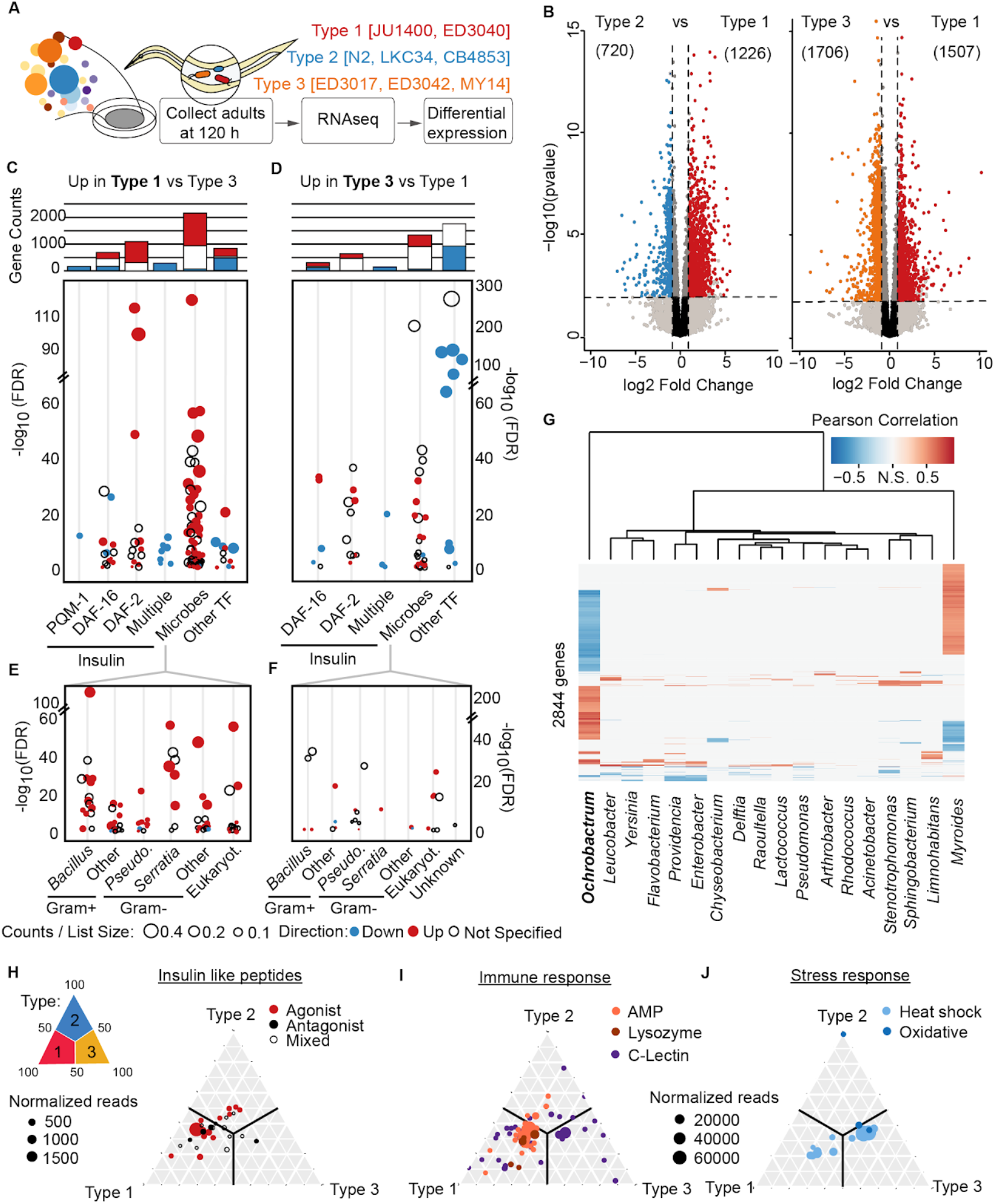
Transcriptional changes in insulin signaling, microbial and stress response genes define microbiome types. **A.** Representative strains from each of the microbiome types grown on BIGbiome to Day 3 adulthood were collected for RNAseq. **B.** Volcano plots displaying genes differentially expressed between Types 1 and 2 and Types 1 and 3. Type 1 vs Type 2; Significantly differentially expressed genes (Benjaminii-Hochberg adjusted p-value < 0.05) are colored red if they are upregulated in Type 1, or log2FC > 1, or colored blue if upregulated in Type 2, or log2FC < −1. Type 1 vs Type 3; Significantly differentially expressed genes (Benjamini-Hochberg adjusted p-value < 0.05) are colored red if they are upregulated in Type 1, or log2FC > 1, or colored orange if upregulated in Type 3, or log2FC < −1. **C-F.** Significant (FDR < 0.05) WormExp enrichments from Type 1 Up gene set (C) and Type 3 Up gene set (D). Barplots represent counts of unique genes for each category. ‘Multiple’ category includes *daf-2;daf-16* double mutants. ‘Microbes’ category subset separated into specific terms in the Type 1 Up set (E) and Type 3 Up set (F). **G.** Heatmap depicting genes that are significantly differentially expressed between microbiome types and significantly correlated (Pearson correlation, Benjamini-Hochberg adjusted p-value < 0.05) with the absolute abundance of at least one BIGbiome member. **H-J.** Ternary plots illustrating the microbiome type enrichment patterns of genes belong to insulin-like peptides (H), immune (I) and stress responses (J). Each dot is an individual gene and dot sizes are proportional to normalized read counts in the transcriptional dataset. Due to a large number in the immune and stress response gene, only genes with significant changes (p< 0.05) in expression between the microbiome types are shown. Only one gene (*ctl-1*) is expressed almost exclusively in Type 2 (in J).

Microbial response pathway genes varied significantly between the strain groups. We found that genes more highly expressed in Type 1 animals were broadly enriched in genes altered in response to a wide array of microbes [60.4% (81/134 datasets) for ‘Type 1 Up’ versus 42.9% (27/63 datasets) for ‘Type 3 Up’; **Figure 3E-F**]. Interestingly, ‘Type 1 Up’ genes overlap with those upregulated upon exposure to pathogens [**Figure 3E**] (e.g., *B. thuringiensis, S. marcescens, E. faecalis, P. aeruginosa* and others ^32–34)^ while ‘Type 3 Up’ genes overlap more with those downregulated upon pathogen exposure [**Figure 3F**]. Though no pathogens are included in the BIGbiome, Type 1 animals seem to be using similar responses to related microbes to exclude most everything but *Ochrobactrum* from the gut. Consistent with this idea, ‘Type 1 Up’ genes overlap with 13 datasets of upregulated genes in response to the pathogen *P. aeruginosa* PA14 [162 genes in total; e.g., 39 genes from ^35^, Q = 1.8e-8]. Under these conditions, the twelve *Pseudomonas* strains in the BIGbiome are excluded from the guts of Type 1 animals [1.2% relative abundance compared to 6.4% and 7.4% for Type 3 and lawns, respectively]. Further, several canonical *C. elegans* immune effectors from multiple pathways ^36^ were expressed more highly in Type 1 animals [**Figure 3I**, **Table S6**]— e.g., *irg-5* [2.81-fold and Pearson=0.75 to *Ochrobactrum*; p38/MAPK and FSHR-1], *lys-5* [2.80-fold; Wnt/β-catenin and HLH-30/TFEB], and *irg-2* [2.9-fold; ZIP-2]. These specific responses are likely to promote *Ochrobactrum* colonization in the process. In contrast, more limited immune pathway expression was observed in Type 3 animals and instead more highly express general stress-related pathways [**Figure 3I-J, Table S6**]— e.g., *gcn-1* [3.2-fold; SKN-1/Nrf2, oxidative stress] and *hsp-6* [2.4-fold; ATFS-1, unfolded protein stress]. Type 3 animals did express a subset of c-type lectins more significantly than the other microbiome types however [**Figure 3I**]. Thus, Type 1 animals appear to utilize a suite of immune pathways in parallel to create the highly selective environment within the gut for *Ochrobactrum* colonization, which are largely absent in Type 3 animals.

### Transcriptional variation in insulin signaling networks distinguish microbiome types

Among the pathways enhanced among the microbiome types, we observed a particular enrichment for insulin signaling. Both ‘Type 1 Up’ and ‘Type 3 Up’ gene sets were highly enriched for DAF-2-and/or DAF-16-dependent genes, though overlap was more extensive in Type 1 animals [47 datasets for ‘Type 1 Up’ and 20 datasets for ‘Type 3 Up’; both high- and low-insulin conditions observed; **Figure 3C-D**]. In addition, the vast majority of the microbially responsive genes identified above are also associated with changes in insulin signaling pathways in the lab strain N2 [1066 genes (80%) in Type 1 versus 464 genes (46%) overlap with insulin signaling datasets; **Table S6, Figure 3C-D**]. Interestingly, the enrichment observed for ‘Type 1 Up’ genes have been associated with both low- and high-insulin signaling conditions in the lab strain N2, which may reflect plasticity in gene expression driven by natural genetic variation in these wild strains.

To more clearly gauge the insulin signaling balance in these animals, we examined expression of the nearly 40 insulin-like peptides (ILP) that compete for binding to DAF-2/IGFR. This mixture of ILPs serves to activate (agonists) or repress (antagonists) downstream insulin signaling pathways to provide phenotypic specificity and coordination of responses across tissues ^37,38^. There was a notable shift in ILP expression between Type 1 and Type 3 animals: 9 of the 40 ILPs were expressed significantly higher in Type 1 strains compared to minimal ILP expression in Type 3 strains [**Figure 3H**]. Type 2 animals expressed intermediate levels and a mix of agonist and antagonist ILPs, consistent with the intermediate expression of insulin pathway genes [**Figure S8A**]. Nearly all of the genes in the canonical insulin signaling pathway, including *daf-2/IGFR, age-1/PI3K, akt-1/AKT, daf-18/PTEN* and *daf-16/FOXO* were expressed higher in Type 3 than Type 1 animals [**Figure S8A**]. This suggests that Type 1 animals are likely responding favorably to the microbiome under high insulin conditions, while Type 3 animals responses are likely less favorable and driven by a scarcity of insulin ligands.

### Insulin signaling pathways drive microbiome composition and its impact on host physiology

We next sought to test directly whether host insulin signaling mediates microbiome selection and its resulting effects on host physiology. To do this, we used RNAi to knock down *daf-2/IGFR* and *daf-16/FOXO* gene expression in representative strains for each microbiome type: Type 1, JU1400; Type 2, N2; and Type 3, ED3017. If high levels of insulin signaling positively select for Type 1 microbial communities, then reducing the activation of these pathways may result in these hosts adopting communities and host physiological attributes that more closely resemble those in Type 3 strains. Indeed, this is what we observed. Knockdowns of *daf-2* resulted in slower development [**Figure S7AB**] and reduced body size in JU1400 [**Figure S7C**] when grown on BIGbiome lawns versus vector controls. Conversely, knockdowns of the transcription factor *daf-16*/FOXO generally accelerated development [Figure S7A,B] and increased animal body size [**Figure S7C**]. In Type 2 animals (N2), we observed lower *Ochrobactrum* colonization in *daf-2* RNAi (P<0.001) and higher in *daf-16* RNAi, as shown via microbiome sequencing and fluorescence quantification of GFP-*Ochrobactrum* [P<0.001, **Figure 4A-C**]. The impact of these knockdowns were most dramatic in Type 1 and 3 strains: *daf-2* RNAi reduced the recruitment of *Ochrobactrum* in Type 1 (JU1400) animals by 30-50% compared to vector controls [P<0.001, **Figure 4A-C**], while *daf-16* RNAi increased *Ochrobactrum* colonization by more than 20-fold in non-selective Type 3 animals (ED3017) [P<0.001, **Figure 4A-C**].

**Figure 4.**
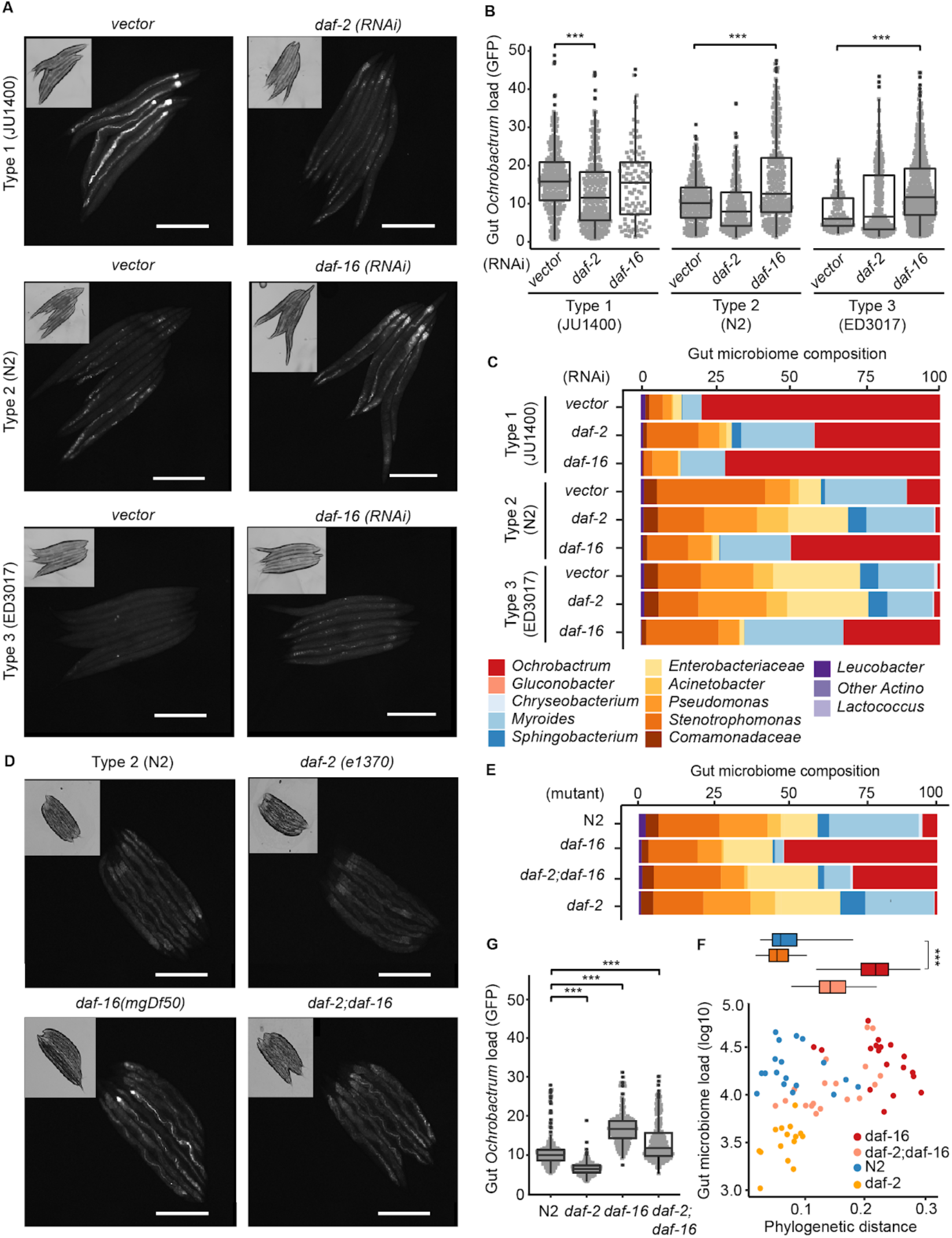
Insulin signaling pathways mediate recruitment of *Ochrobactrum*. **A-C.** *Ochrobactrum* colonization in JU1400(Type 1) decreased with *daf-2*(RNAi) and increased in N2(Type 2) and ED3017(Type 3) with *daf-16*(RNAi). Similar trends are shown in representative images of Day 3 adults grown on BIGbiome with GFP-*Ochrobactrum* (A, Bar = 500 μm), GFP signal per individual animal (B, JU1400(vector): n=290, JU1400(*daf-2*): n=274, JU1400(*daf-16*): n=97, N2(vector): n=233, N2(*daf-2*): n=202, N2(*daf-16*): n=79, ED3017(vector): n=103, ED3017(*daf-2*): n=247, ED3017(*daf-16*): n=292), and bulk gut microbiome sequence of the corresponding population (C). **D,G.** *Ochrobactrum* colonization decreased in *daf-2*(e1370), increased in *daf-16(mgDf50)* and by a lesser extent in *daf-2(*e1370*);daf-16(mgDf50)* mutants. Similar trends are shown in representative images of Day 3 adults grown on BIGbiome with GFP-*Ochrobactrum* (D, Bar = 500 μm), bulk gut microbiome sequence of the corresponding population (E), and GFP signal per individual animal (G, N2: n=216, *daf-2*: n=198, *daf-16*: n=184, *daf-2;daf-16*: n=207). **F**. N2 and insulin signaling mutants *daf-2*, *daf-16*, *daf-2;daf-16* host distinct microbiome types based on gut microbiome load per animal (y-axis) and phylogenetic distances to BIGbiome lawn (x-axis). Inset: Box-whisker plot of phylogenetic distances to BIGbiome for the three microbiome types. (B,G) n=individual animals; P-values were generated from one-way ANOVA, followed by and post hoc Tukey test with 95% confidence level and adjusted for multiple comparisons (***p<0.001, **p<0.01, *p<0.05).

Since Type 2 animals are intermediate in their selectivity for *Ochrobactrum*, we next sought to test whether Type 2 insulin signaling mutants change their microbiome type. Loss-of-function mutants of the insulin peptide receptor *daf-2*(e1370) in the lab strain (N2) mimic low insulin levels. On BIGbiome lawns, *daf-2*/IGFR mutants exhibited Type 3-like developmental delays and smaller body size compared to wild type animals [P<0.001; **Figure S7D-E**]. The *daf-2*/IGFR mutants had lower *Ochrobactrum* colonization at the population level by microbiome sequencing [P<0.001; **Figure 4E**] and at individual level by fluorescence quantification of GFP-*Ochrobactrum* [P<0.001; **Figure 4G**]. Conversely, loss-of-function mutants of the downstream transcription factor *daf-16*(mgDf50) developed much faster, had larger body sizes in early adulthood [P<0.001; **Figure S7D-E**], and had greater *Ochrobactrum* colonization [P<0.001, **Figure 4D-G**]. Finally, double-mutants of *daf-2;daf-16* increased *Ochrobactrum* colonization by one-third compared to *daf-16* mutants [P<0.001; **Figure 4D-G**], suggesting other potential regulators may be acting in the low insulin signaling conditions to suppress *Ochrobactrum* colonization. Together, these data indicate that under low insulin signaling, DAF-16 regulated processes either limit *Ochrobactrum* colonization or fail to effectively exclude other microbiome members.

To test the generalizability of these responses, we then expanded our RNAi analyses to both additional representative strains [ED3042 (Type 3), CB4856 (Type 1) and N2 (Type 2)] and additional genes in the canonical insulin signaling pathway. We observed that RNAi-mediated knockdown genes that activate the insulin signaling like *daf-2/IGFR*, *age-1/PI3K* and *akt-1/AKT* all reduced *Ochrobactrum* colonization levels in Type 1 and 2 animals [**Figure S8C**]. Conversely, knockdowns of *akt-2/AKT* and pathway suppressor *daf-18/PTEN* increased *Ochrobactrum* colonization [**Figure S8C**] in Type 2 and 3 animals. Further, knocking down each of the 20 insulin-like peptides in the intestine of Type 2 (N2) animals, increased *Ochrobactrum* colonization for 6 of the 9 antagonist ILPs [*ins-1, 11, -18, -21, -24, - 31*; P<0.001, **Figure S8B**]. Together, these data indicate that subsets of canonical insulin signaling pathways are utilized across wild *C. elegans* strains to regulate microbiome composition and, in turn, impact host physiology and growth.

### Interplay of downstream insulin signaling transcription factors drives microbiome regulation

We next sought to determine what genes in the insulin signaling regulons influence microbiome composition. To achieve this, we examined two mutually exclusive transcription factors known to orchestrate insulin signaling in *C. elegans*, DAF-16/FOXO and PQM-1/SALL2. PQM-1 has also previously been associated with regulation of both development and immunity into adulthood ^39,40^. To directly test its role in regulation of the microbiome we knocked down *pqm-1* by RNAi in representative strains of each microbial community type: JU1400 (Type 1), N2 (Type 2) and ED3017 (Type 3). We observed significantly delayed development and reduced body sizes on BIGbiome in Type 1 animals [P = 0.02 and 0.004, respectively; **Figure 5A-B**]. Consistent with the effect on development, we observed a decrease of *Ochrobactrum* colonization after *pqm-1* knockdown by RNAi [P<0.001; **Figure 5E-F**]. Knockdowns of *pqm-1* in Type 2 (N2) and Type 3 (ED3017) animals showed similar but non-significant decreases in developmental rates [**Figure S9A-B**], and the already low levels of *Ochrobactrum* colonization were decreased to the limit of detection for both strains [**Figure S9C**].

**Figure 5.**
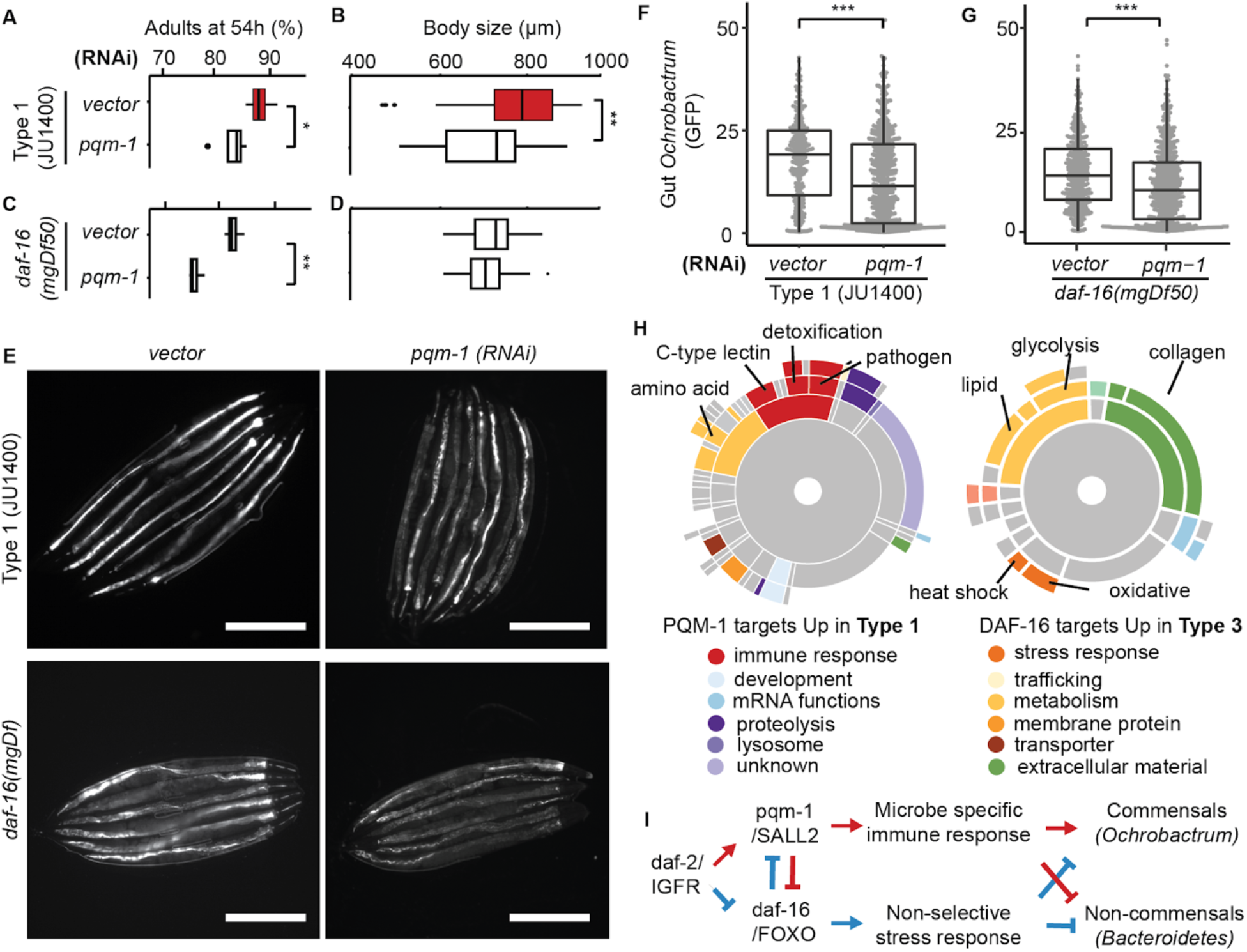
PQM-1 regulates microbiome impact on host physiology and recruitment of *Ochrobactrum* to the gut microbiome. **A.** Box-whisker plot of adult percentage of vector (n=4) and *pqm-1* (n=4) RNAi knockdown mutants in Type 1 JU1400 background at 54 h post L1 stage. **B.** Box-whisker plot of body size of vector (n=66) and *pqm-1* (n=85) RNAi knockdown mutants in Type 1 JU1400 background at 48 h post L1 stage. **C.** Box-whisker plot of adult percentage of vector (n=4) and *pqm-1* (n=4) RNAi knockdown mutants in *daf-16(mgDf50)* background at 54 h post L1 stage. **D.** Box-whisker plot of body size of vector (n=136) and *pqm-1* RNAi knockdown mutants (n=179) in *daf-16(mgDf50)* background at 48 h post L1 stage. **E-G.** *Ochrobactrum* colonization in JU1400(Type 1) and *daf-16(-)* decreased with *pqm-1*(RNAi). Similar trends are shown in representative images of day 3 adults grown on BIGbiome with GFP-*Ochrobactrum* (**E**, Bar = 500 μm) and GFP signal per individual animal (**F**,**G**). (**B**,**D**) n represents the number of independent worm populations. (**C**,**E**) n represents the number of individual animals quantified by microscopic images. (**G**,**H**) n represents the number of individual animals quantified by Biosorter. P-values were generated from student’s t-test (***p<0.001, **p<0.01,*p<0.05). **H.** Sunburst plot illustrating significantly enriched (WormCat-reported padj < 0.05) WormCat subcategories from Class II targets upregulated in Type 1 strains and Class I targets upregulated in Type 3 strains. **I.** Schematic diagram of insulin signaling targets drives *Ochrobactrum* colonization (Type 1, red arrows; Type 3, blue arrows).

To further dissect the genetic interaction of *pqm-1* with *daf-2* and *daf-16*, we knocked down *pqm-1* in *daf-2*(e1370) mutants by RNAi and observed significantly slower developmental rates [40% less, P<0.001; **Figure S9D**] and reduced body sizes [13%, P<0.001; **Figure S9E**], but *Ochrobactrum* colonization remained similar to the empty vector at a low level. RNAi knockdown of *pqm-1* in *daf-16*(mgDf50) mutants also delayed development by 10% [P<0.001; **Figure 5C**] and significantly lowered *Ochrobactrum* colonization compared to the empty vector [**Figure 5E,G**]. These data suggest that *pqm-1* promotes *Ochrobactrum* colonization independent of *daf-16*.

Finally, we examined the transcriptional networks themselves based on promoter binding elements for each of these transcription factors. DAF-16 activates genes that contain a DAF-16 binding element (DBE; Class I) under stressful or low insulin conditions, while PQM-1 activates genes under favorable or high insulin conditions containing the DAF-16 associated element (DAE; Class II) ^39^. Analysis of the transcriptional datasets identified 170 (10.2%) Class I and 219 (12.6%) Class II genes that were differentially regulated between Types 1 and 3. WormCat analyses of these genes highlighted two very different responses in Type 1 and Type 3 animals. The Class II genes from the ‘Type 1 Up’ set [**Figure 5H**] are enriched for multiple detoxification and immune responses against pathogens, including cytochrome P450 genes (*cyp-13A3, cyp-32A1, cyp-25A1*), which can metabolize toxic compounds, and c-type lectins (*clec-57, clec-49, clec-204*), which are involved in antimicrobial immunity ^51^. In contrast, the Class I genes from the ‘Type 3 Up’ set are enriched for general oxidative and heat stress responses rather than pathogen specific responses. Metabolism categories also differed between Types 1 and 3, with Type 3 enrichment for glycolysis, lipid (fatty acid and phospholipid), short chain dehydrogenase, and carbohydrates. Favorable insulin signaling may therefore promote the selection of more specialized microbial communities via regulation of immune and xenobiotic response genes that may help establish a selective environment for *Ochrobactrum* to colonize the gut. Our results suggest higher insulin signaling activates PQM-1 to promote microbial specific immune response in commensal selection from the environment, while lower insulin signaling levels drive DAF-16 mediated broad stress responses that suppress microbiome selection [**Figure 5I**].

## DISCUSSION

### Regulation of the gut microbiome is driven by host genetics in *C. elegans*

*C. elegans* flourish in natural habitats of rotten fruit and plant matter, an environment with abundant and diverse microbes. They rapidly respond to environmental fluctuation and adjust growth, defense and reproduction strategies to ensure their success in the wild. To learn more about the genetic circuits in microbiome response from this widely used model organism, we reunite wild *C. elegans* with microbial consortia isolated from their natural habitat. Although grown on the same microbiome mixture BIGbiome, 38 *C. elegans* strains established distinct gut microbiome types in adulthood, likely driven by their natural genetic variation. Wild *C. elegans* with stronger recruitment of *Ochrobactrum*, a commensal member of their core microbiome in nature, demonstrated faster growth and development. Transcriptomic analysis suggested host insulin signaling was driving the *Ochrobactrum* dominant gut microbiome. We used RNAi knockdown and mutants to confirm that IIS modulates the *Ochrobactrum*-driven microbiome type variation through downstream transcription factors of DAF-16/FOXO and PQM-1/SALL2.

The influences of highly conserved insulin signaling pathways are found in nearly all aspects of animal physiology, including development, fertility, stress resistance and longevity ^41–43^. Evidence of gut microbes engaged in IIS are found in fruit fly, zebrafish, mouse, and human ^1,44,45^, where low insulin signaling is typically associated with benefits to host physiology (longer life, increased pathogen resistance, and the like). Our findings underscore a distinct role for insulin signaling in establishing both a selective environment for microbiome enrichment.

*C. elegans* grew and developed faster on BIGbiome than on OP50, particularly for microbiome Type 1 strains that selected commensal *Ochrobactrum*. It is possible that commensal microbes in BIGbiome stimulated host IIS that accelerated their growth and development. In return, elevated IIS created a suitable gut environment that allowed more commensal microbes to colonize. The microbe-IIS-microbe interactions formed a positive feedback loop to establish an *Ochrobactrum* dominant microbiome type in adulthood. On the other hand, strains that failed to establish this positive feedback interaction then activated broad-spectrum stress responses that limit microbiome colonization, but meanwhile abolished the ability to select commensals from the environment.

### Insulin signaling mediated commensal selection (Type 1)

Our study further suggested how the IIS mediated commensal selection unfolded in *C. elegans*. From upstream of the signaling cascade, insulin-like peptides in the intestine mediate the microbe-host communications to balance energy expenditures in growth, reproduction, and defense. Indeed, ILPs are increasingly expressed in the intestine with age ^37^, suggesting their critical roles in decision-making during adulthood. 40 insulin like peptides (ILP) compete for binding to DAF-2/IGFR, functioning either to activate (agonists) or repress (antagonists) downstream signaling ^46,47^; *C. elegans* uses this network of ILPs to provide phenotypic specificity and coordination across tissues ^38^. INS-7 is an agonist of DAF-2/IGFR and is regulated by DAF-2/IGFR and DAF-16/FOXO in the intestine to provide positive feedback regulation in coordination of animal physiology across tissues ^37^. Thus, we hypothesize that the observed high expression of *ins-7* and other ILPs in Type 1 keeps DAF-16 activity low to prevent overstimulation of the immune system, or indiscriminate microbial response leading to commensal exclusion. On the other hand, multiple studies found pathogen infections induced antagonistic *ins-11* expression through HLH-30 dependent p38 MAPK pathway ^48^. We confirmed by RNAi knockdown in N2 that *ins-11* from the intestine suppressed gut *Ochrobactrum* colonization. It is possible that more pathogenic colonizers from the BIGbiome can suppress host IIS by stimulating antagonistic ILPs production, thus reducing host selection of commensals, forming the microbiome types 2 and 3 we observed.

IIS regulated PQM-1 drives downstream targets with GATA promoter motif ^49^, likely contributing to gut microbiome selection. In the *Ochrobactrum*-dominant microbiome type 1, up-regulated PQM-1 targets are enriched in host immune response genes, including C-type lectin and antimicrobial peptide ^50^. C-type lectins are known to recognize microbial molecular patterns, implying their roles in bacterial specific immunity ^51^. Antimicrobial peptides were also enriched as up-regulated PQM-1 targets, among them saposin genes like *spp-2* and *-5* that were known to be induced by *Ochrobactrum* colonization ^26^. In addition, the *Ochrobactrum*-dominant microbiome type is associated with elevated xenobiotic response gene families like CYP, GST, and UGT. These enzymes can detoxify microbial products and act as a sink of reactive oxygen species (ROS), thus reducing oxidative stress for cellular protection and maintenance. Taken together, these responses may create a gut environment that favors *Ochrobactrum* and gut microbiome selection.

### Trade-off in loss of insulin signaling pathway (Type 3)

Reduced IIS in strains with non-selective microbiome type resulted in stronger oxidative stress, as shown in higher expression of catalase genes, as well as mitochondria and ER stress markers like *hsp-4* and *-6*, mimic stress related phenotypes that were observed in the long-lived *daf-2* mutant ^43,52^. Other signatures of *daf-2* mutants include shift of lipid metabolism and reduced brood size. Similarly, microbiome Type 3 strains increased the expression of mitochondrial β-oxidation genes like *acdh-2* and glycogen synthesis genes like *gsy-1*, indicating a switch from lipid metabolism to carbohydrate storage. In addition, transcription factors (*lin-11*, *lin-13b*, *mep-1*) that negatively regulated reproduction were highly expressed in non-selective Type 3 strains, suggesting a reduced investment in reproduction. Interestingly, increased fertility was observed in *C. elegans* colonized by *Ochrobactrum*, driven by genes with enriched GATA motifs ^26^. Therefore, it is possible that PQM-1 activates these *Ochrobactrum*-responsive genes in adulthood, linking commensal recruitment to evolutionary benefits in host development and reproduction. As the worm ages, reduced insulin signaling during adulthood activates DAF-16 dependent immune response to defense against microbes, compensating for the immune-senescence in other protective pathways like the MAP kinase ^53^. Although reduced insulin signaling provides benefit to the host in pathogen resistance and lifespan extension, the trade-off in adulthood might be loss of commensal colonization and reduced reproduction.

### Broader signaling networks in regulation of the microbiome

Acting in the same direction of IIS, TGF-β signaling was also up-regulated in the *Ochrobactrum* dominant microbiome Type 1 animals, likely the result of extensive crosstalk between the two pathways ^54^. TGF-β signaling from neurons and epidermis can activate ILPs secretion that feed into IIS to modulate DAF-16 activities in the intestine ^55,56^. *C. elegans* TGF-β mutants *dbl-1* were highly colonized by *Enterobacter* with enhanced pathogenicity when grown on a synthetic natural microbiome, and though these genes remain unchanged in these studies, it still highlights the importance of this pathway in microbiome regulation ^57^.

Similar to DAF-16, transcription factor SKN-1 was also enriched in the intestine and acts downstream of IIS as an AKT-1 phosphorylation target ^58^. Under reduced IIS in the microbiome type 3 strains, SKN-1 likely synergized with DAF-16 to induce oxidative and heat shock stress that suppress microbiome selection and colonization, which explained why *daf-2;daf-16* double mutant only partial reduced *Ochrobactrum* colonization compare to *daf-2* mutant. Interestingly, the longevity effect of SKN-1 was dependent on type of *E. coli* strains, suggesting the pathway is under the influence of microbial content ^59^. SKN-1 may be also responsible for higher collagen genes in microbiome type 3, as these extracellular matrix (ECM) genes were known to be up-regulated by SKN-1 and play critical roles in pathogen defense as weakened cuticles were associated with increased susceptibility to *Microbacterium nematophilum* infection ^60,61^.

### Prospectus

Animals have partnered with microbes throughout evolution to extend their genetic repertoire and metabolic capacity ^62^. The partnership is now deeply imprinted in animal physiology that disruption of this commensal relationship will compromise animal health. Multiple animal models were established to decode this evolutionary conserved program of interactions. Here, we presented a genetically tractable platform that integrates natural microbiome with rich molecular tools in *C. elegans*. Our results demonstrated that natural variation in IIS drives microbiome selection, suggesting that the regulon of *daf-16* and *pqm-1* played major roles in the *C. elegans* microbiome types formation. Further evidence from posttranslational modification and nuclear localization of these transcription factors are needed to confirm tissue-specific roles in responding to microbes. Additionally, long-term life history readouts of microbiome selection like reproduction and lifespan remain to be investigated. From the microbial side, increasing genomic information from natural microbiome ^20^ will undoubtedly help the discovery of commensal metabolites that engage in host pathways like IIS. Moreover, over 40% of microbiome type enriched genes we identified belong to understudied genes without known function. It is possible that interactions with the microbiome from native habitats induced host responses that were absent before, presenting additional opportunities to reveal novel functions and untangle the intertwined network between microbiome and host.

## ACKNOWLEDGEMENTS

This work was supported by NIH grants DP2DK116645 (to B.S.S). This project was supported by the Cytometry and Cell Sorting Core at Baylor College of Medicine with funding from the CPRIT Core Facility Support Award (CPRIT-RP180672), the NIH (S10 OD025251, CA125123, and RR024574) and the assistance of Joel M. Sederstrom, plus an instrumentation grant for the Biosorter NIH grant (S10 OD025251). Some strains were provided by the CGC, which is funded by NIH Office of Research Infrastructure Programs (P40 OD010440). Emily Troemel generously provided *Ochrobactrum* BH3 and isogenic GFP expressing strains. We also thank Gretchen Diehl Lab, Joseph Hyser Lab, Rachel Arey Lab, Meng Wang Lab and Houston Area Worm Group and Center for Metagenomics and Microbiome Research labs for helpful advice at various stages of this project, plus Diehl and Estes labs for sharing key equipment and the CMMR core facility for microbiome sequencing and Rachel Arey for helpful comments on the manuscript.

## AUTHOR CONTRIBUTIONS

Conceptualization, F.Z., J.L.W., B.S.S.; Methodology Development, F.Z., J.L.W., A.A., C.A.A., M.L.C.; Software Programming, F.Z., J.L.W., A.A.; Validation, F.Z., A.A., A.S.K.; Formal Analysis, F.Z., J.L.W., A.A., C.H., B.S.S.; Investigation, F.Z., J.L.W., C.H.; Resources, M.-A.F., M.L.C., D.V.V., C.A.A.; Data Curation, F.Z., J.L.W., A.A.; Writing – Original Draft, F.Z., J.L.W., B.S.S.; Writing – Review & Editing, F.Z., J.L.W., B.S.S., M.-A.F., A.A., C.A.A., M.L.C.; Visualization, F.Z., J.L.W.; Supervision, F.Z., J.L.W., B.S.S.; and Project Administration and Funding Acquisition, B.S.S.

## MATERIALS AND METHODS

### RESOURCE AVAILABILITY

#### Lead Contact

Further information and requests for resources and reagents should be directed to and will be fulfilled by the Lead Contact, Buck Samuel.

#### Materials Availability

All microbial strains and other materials used in these studies are available upon request.

#### Data and Code Availability

All datasets have been included as raw data [**Table S7**]. Sequencing based datasets have been deposited at NCBI Sequence Read Archive database (Bioproject PRJNA540192) with the following sample accession numbers for RNAseq reads (SAMN13050735-13050742) and microbiome sequencing reads (SAMN13068200-13068238, 13071563-13071602, 16597785-16597833, 16611296-16611371, 17054579-17054627). All code used in the analysis of datasets is included in supplemental information as an archive [**File 1**].

### EXPERIMENTAL MODEL AND SUBJECT DETAILS

#### Maintenance of Caenorhabditis elegans strains

*Caenorhabditis elegans* strains utilized in this study can be obtained from the *Caenorhabditis* Genetics Center (CGC), including N2-Bristol, CB1370 [*daf-2*(e1370)], GR1307 [*daf-16*(mgDf53)], HT1890 [*daf-2*(e1370);*daf-16*(mgDf53)] and several natural isolates: AB1, AB3, CB4853, CB4854, CB4856, ED3017, ED3021, ED3040, ED3042, ED3052, ED3072, GXW0001, JU1088, JU1171, JU1218, JU1400, JU1401, JU1652, JU258, JU263, JU300, JU312, JU322, JU323, JU360, JU361, JU397, JU533, JU642, JU775, KR314, LKC34, MY1, MY14, MY16, MY2, and PX174 [**Table S2**]. The intestinal RNAi strain JM45 (*rde-1(ne219); Is[Pges-1::RDE-1::unc54 3*′*UTR; Pmyo2::RFP3]*) was a gift from Dr. Meng Wang. All *C. elegans* strains were grown and maintained on nematode growth media (NGM; Research Products International) seeded with *Escherichia coli* strain OP50 at 20°C. *E. coli* OP50 and HT115 RNAi strains can be requested from the CGC.

#### Preparation of *C. elegans* populations

Prior to each experiment, worm populations were rendered ‘germ-free’ and synchronized to L1 stage ^63^ by treating gravid hermaphrodites with bleach solution (mixture of Clorox bleach and 5M NaOH in 2:1 volume ratio), followed by multiple washes with M9 buffer ^63^ to remove bleach solution. Germ-free L1s were then allowed to hatch and synchronize in sterile M9 buffer 15-18 hours rotating at 20°C.

#### Preparation of microbiome mixtures

All microbial strains used were originally isolated from *C. elegans* natural isolates or habitats [**Table S1**] and stored at −80°C as glycerol stocks ^16^. *Ochrobactrum pituitosum* BH3 and an isogenic strain expressing GFP [Tn7 insertion of GFP on the chromosome; ^64^] were generous gifts from Dr. Emily Troemel. JUb strains were originally isolated by Dr. Marie-Anne Félix.

To begin all experiments, we stamped out fresh cultures from glycerol stocks onto a rectangular LB plate, then incubated overnight at 28°C. The colonies on the plate were then used to inoculate a 1 ml 96 deep well plate (Axygen) filled with 300 μl lysogeny broth (10g Tryptone, 5g yeast extract, 10g NaCl in 1L distilled water adjust to pH=7.5) in each well. After overnight growth (14-16 h) at 28°C and 250 rpm shaking, bacterial cells were pelleted down by centrifuge at 4000 x g for 10 min. Supernatants were discarded and replaced with 200 μL sterile M9 buffer in each well. Pellets were then fully resuspended by pipetting then transferred to a clear bottom 96 well plate (Costar, Corning). Growth of each microbe was assessed by measurement of optical density (OD) readings at 600nm using a Multiskan FC Microplate Photometer (Thermo Scientific). Bacterial density in each well in the parent plate was then normalized individually to an OD_600_ of 1.0 using sterile filtered M9 buffer. BIGbiome001 master mixes (referred to as ‘BIGbiome’ throughout) were created by combining equal volumes of each bacterial strain, which was then used to seed (30 μL) Nematode Growth Medium (NGM) agar in 12 well plates (Costar, Corning). Seeded plates were grown overnight at 20°C (80% humidity) before use.

### METHOD DETAILS

#### Measurement of gut microbiome colonization in *C. elegans*

Existing methods that use surface sterilization with antibiotics, pestle-based disruption of animals and enumeration of bacterial colonies on agar plates^21^, though robust, were optimized for determination of bacterial densities of an individual strain or small set bacteria of interest rather than communities. Discrimination of bacteria by colony morphologies is similarly intractable within complex communities. We addressed these challenges by: (i) replacing antibiotic treatment, which is ineffective in a large community that contains variable antibiotic resistance profiles, with a more consistent dilute bleach treatment to kill surface associated microbes; and (ii) replacing the mortar-and-pestle with bead-based, multi-well format disruption of *C. elegans* to release gut microbes into solution. Further, to quantify live bacteria in the gut, we also adapted a liquid-based CFU quantification method to remove the need for laborious colony counting on plates.

##### Creation of standard curves for CFU estimations

Overnight grown BIGbiome lawn was sampled and resuspended in M9 buffer. The mixture was subjected to a serial dilution from 10^−1^ to 10^−6^. The number of live bacteria from the dilution series were determined by counting CFU from 10 μL of each dilution onto a LB plate. The same dilution was inoculated into a 96 well flat bottom plate containing 100 μL LB medium in each well. The plate was incubated at 28 °C and bacterial growth curve in each dilution was recorded by measuring OD_600_ every 15 min for 18 h. Within the range of linear portion of growth, OD_600_ equal to 0.2 was used as a threshold to interpolate the corresponding growth time, designated as CGT ^65^. Exponential regression between CFU number and CGT (R² = 0.99) was used to infer the CFU number from sample CGT at OD_600_ threshold of 0.2. Regression derived trendline equation was applied: total bacterial cells = (8E+11)*e^(−1.114 * CGT)^.

##### Collection, surface sterilization and lysis of animals

Around 100 L1 animals were seeded in duplicate on the BIGbiome lawn at 20°C with 80% humidity. Worm populations were assayed at 48 h and 120 h post seeding. On sampling day, worms were washed from a bacterial lawn with 600 μL of M9 buffer (0.01% triton X-100) to a sterilized 2 ml 96-well deep plate (Axygen). The deep well plate was centrifuged at 300 g for 1 minute to pellet down worms, bacteria in the liquid were removed by an aspirating manifold (VP1171A, V&P scientific). These washing steps were repeated 5 times with M9 buffer (0.01% triton X). 100 μL of 10 mM levamisole in M9 buffer (0.01% triton X) was then added to paralyze the worms for 5 min. Then 200μl of 4% bleach solution (diluted from of Clorox bleach and 5M NaOH in 2:1 mixture) in M9 treatment for 2 min, further eliminate residual bacteria in liquid and on worm cuticle. 2 more washing with M9 buffer (0.01% triton X) was done to remove bleach and levamisole solution. After the last wash, An aliquot of liquid volume from each well was transferred to a new flat bottom 96 well plate (Costar 3370, Corning) for bright field imaging under a Nikon TiE Inverted Microscope. Generated images were used to estimate the number and size of adult animals in each well. An aliquot of supernatant from the imaging plate was taken as a negative control to assess background residual live bacteria before host lysis. The remaining worms were then lysed by adding 1.0 mm sterilized garnet beads (Biospect) in a Mixer Mill (Restch) at 25 Hz for 5 min to release live bacteria into solution.

##### Quantification of bacterial densities using growth curve estimations

Worm lysates were diluted 10 fold with M9 buffer to reduce debris, and 20 μL of the lysate dilution was inoculated into a 96 well flat bottom plate with 100 μL LB medium. The plate was incubated at 28°C for 18 h. OD600 values were recorded every 15 min to generate bacterial growth curves for each well. Threshold growth time (CGT) at OD600 equal to 0.2 was derived from the corresponding growth curve. Total bacterial cells in each well were calculated based on the BIGbiome equation with corresponding CGT number. Colonization level per animal was then calculated using the following formula:

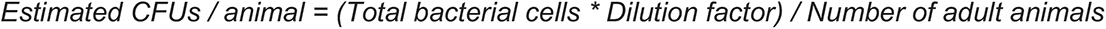

#### Measurement of gut microbiome composition in *C. elegans*

##### Collection and lysis of animals

Worm lysate from the previous step by centrifuged at 4000 x g for 10 min. extraction, a freeze-thaw process in −80°C freezer overnight was first applied, then 0.1 mm sterile zirconia/silica beads (BioSpec products) were added (enough to cover well bottom), bead-beating in Mixer Mill (Restch) at 25 Hz for 5 min to disrupt bacterial cells. Immediately followed by enzymatic treatment of 1 mg/mL proteinase K (NEB) at 60°C for 60 min, then 95°C for 15 min to deactivate the proteinase K. After the treatment, samples were centrifuged at 4000 g for 10 min to pellet down cellular fractions.

##### Amplicon library construction and sequencing

Supernatant from lysate was transferred to a clean 96 well PCR plate as DNA template. 16S rRNA gene primer set (515F/806R) targeting variable region 4 in bacteria ^66^. Barcode information was added to the reverse primer 806r. Amplicons for each library were normalized based on the PCR product quantified by image processing package in Fiji, then pooled into a single tube for Illumina MiSeq. A detailed protocol for high throughput colonization assay can be found on protocols.io (DOI: dx.doi.org/10.17504/protocols.io.rtzd6p6).

##### Analysis of gut microbiome composition

Fastq files for each library were split by barcode and quality trimmed in the QIIME software package (v1.9.0) ^67^ with an average quality score of 30. Chimeras were removed by usearch61 and Greengenes 13.8 database. Resulting fasta files were imported to Deblur ^68^ with default parameters with all sequences trimmed to 250 bp and positive filter based on 16S rRNA sequences of the 63 strains in the core microbiome. A phylogenetic tree with all Amplicon Sequence Variant (ASV) detected was generated using maximum likelihood method in Mega7 with default parameters. Diversity indices were computed in QIIME using core_diversity_analyses.py with default parameters and rarefied to 3,000 sequences. Alpha diversity was determined using Faith’s phylogenetic diversity and Beta-diversity (between samples) distance matrices were computed within QIIME using default parameters. Phylogenetic-based weighted UniFrac metric was used to compare compositional overlap between worm microbiomes and BIGbiome lawns; the weighted UniFrac metric refers to the degree of overlap in two communities as a function of taxa abundance and shared branches on a combined phylogenetic tree ^69^. Large ‘distances’ indicate less overlap and distinct community compositions. A detailed working pipeline can be found in supplemental files [**File 1**].

#### GWAS analyses of genetic associations with gut microbiome abundance

The *Caenorhabditis elegans* Natural Diversity Resource (CeNDR) was used to perform GWAS^70^ using the EMMA algorithm via the rrBLUP package ^71,72^. The EMMA algorithm used within CeNDR takes into account prevalent linkage disequilibrium observed in *C. elegans* ^73^. The gut microbiome taxa abundance values and *C. elegans* strain names were used as input for GWAS. The CeNDR version used was 1.2.9, with data release 20180527 and cegwas version 1.01. Version WS263 of the worm genome was used in this data release. Representative strains for isotypes with more than one strain tested were randomly selected prior GWAS analyses in CeNDR.

#### RNAi knockdown of *C. elegans* genes

L1 animals were grown on NGM plates with 25 μg/ml carbenicillin and 1 mM IPTG and seeded with 30 μL (OD=1) of *E. coli* HT115 expressing dsRNA to *C. elegans* target genes. To separate exposures to *E. coli* and BIGbiome communities, RNAi treated gravid adults were treated with bleach solution to generate synchronized L1 progeny. Around 100 L1 animals (RNAi F1s) were transferred to NGM plates with BIGbiome lawn to assess gut microbiome colonization and composition after 120 hrs. Previous studies have shown that progeny typically maintain the RNAi-mediated silencing for at least one generation ^74^. Natural variation in RNAi effectiveness in wild strains of *C. elegans* was also assessed by measuring adult body size following *dpy-13* RNAi knockdowns, and no significant differences were observed (JU1400(vector): 1360±154 μm n=16, JU1400(*dpy-13*): 632±156 μm n=15, N2(vector): 1373±199 μm n=30, N2(*dpy-13*): 655±185 μm n=25, ED3017(vector): 1396±236 μm n=25, ED3017(*dpy-13*): 687±173 μm n=22, TableS7.16).

#### Transcriptional profiling of *C. elegans* animals

##### RNA isolation, library preparation and sequencing

*C. elegans* strains [CB4853, ED3017, ED3040, ED3042, JU258, JU775, JU300, JU1400, LKC34, MY14, and N2] for RNAseq were grown on BIGbiome in triplicate for 120 hrs at 20°C. Animals were then washed off plates using M9 buffer (plus 0.01% triton X-100), and progeny were removed by filtering through a sterile 40 μm Nylon mesh (Fisher Scientific). Approximately 500 adult worms were aliquoted in 1.5 mL Eppendorf tubes and placed on ice for 1 min to settle animals, then combined with 200µL of Trizol and 10-20 1.0mm garnet beads. Animals were lysed using a Mixer-mill (Restch) at 25 Hz for 5 min and then incubated at 4°C for 5 min. 200 μL of chloroform was then added to each tube and vortexed for 30 s to mix, and allowed to incubate at room temperature for 3 minutes. Debris was removed by centrifugation at 13,800 x g for 15 minutes at 4°C, and supernatants (∼200 μL) were transferred into RNase-free Eppendorf tubes and stored at −80 °C until extraction. Frozen supernatants were thawed at room temperature and loaded to a KingFisher flex purification system (Thermo Scientific) for automatic RNA processing using MirVANA total RNA kit (Thermo Scientific) following the manufacturer’s protocol. Purified RNAs were stored at −20°C in elution buffer until use. Aliquots of RNA (0.5-2 μg) were used for creation of RNA sequencing libraries and sequenced by Illumina HiSeq4000 (paired end 150bp reads; QuickBiology).

##### RNAseq processing and analysis

An average of 19,638,470 reads were obtained from each dataset, and samples with less than 2 replicates were not utilized in analyses. RNAseq result quality was examined using FASTQC (https://www.bioinformatics.babraham.ac.uk/projects/fastqc/; ^75^), and reads were filtered and trimmed using bbmap (https://jgi.doe.gov/data-and-tools/bbtools/bb-tools-user-guide/bbmap-guide/) and bbduk, respectively (https://jgi.doe.gov/data-and-tools/bbtools/bb-tools-user-guide/bbduk-guide/; https://sourceforge.net/projects/bbmap/). Reads that did not map to the *C. elegans* genome (build WBcel235) with high quality were removed from the analysis [3.8-5.6% of reads for each dataset]. This was based on an internal evolutionary probability model score ‘minid’, which was set to 0.92, and described in more detail in the bb tools user guide referenced above. Acceptable reads were trimmed using bbduk with the following parameters: ktrim=r, k-23, mink=11, and hdist=1. These and other parameters are described in detail in the bb tools user guide referenced above. Filtered and mapped reads (average 19,586,505 per dataset) were pseudoaligned to the WBcel235 genome assembly using kallisto [https://pachterlab.github.io/kallisto/; ^76^] with default settings [89.0-93.3% aligned for each dataset]. DESeq2 was used to estimate differential expression in an R workspace [https://bioconductor.org/packages/release/bioc/html/DESeq2.html; ^77^]. Briefly, DESeq2 models raw counts, normalizes to library depth, estimates and shrinks gene-wise dispersions, and fits a negative binomial model to estimate differential expression based on a Likelihood Ratio Test. Genes with an adjusted p-value of 0.05 or smaller and expressional change greater than two-fold were considered differentially expressed and used in further analyses.

##### Gene set enrichment analyses

WormExp [https://wormexp.zoologie.uni-kiel.de/wormexp/; ^31^] was used for gene set enrichment analyses compared to a comprehensive database of over 1700 curated gene expression datasets in *C. elegans*. Significance for enrichment scores are calculated using the method developed for the program EASE ^78^ and reported as uncorrected p-value, Bonferroni-corrected p-value, and False Discovery Rate. Terms were considered significant if the WormExp-reported FDR score was less than 0.05. WormCat [http://wormcat.com; ^29^] is a similar nematode-specific enrichment analysis and visualization tool that allows for easy categorization and interpretation of datasets based on gene ontology (GO) terms. WormCat is designed for identification of gene sets that are coexpressed or cofunctioning, allowing for drilled-down analysis of specific pathways. Significance scores are reported as Fisher’s exact test p-values. Terms were considered significant if the WormCat-reported P-Value score was less than 0.05.

#### Quantification of animal body size and *Ochrobactrum* gut colonization

##### Microscopy-based quantification

GFP expressing *Ochrobactrum* were used to visualize the colonization of this bacterium. Brightfield and fluorescent images taken by a Nikon TiE Inverted Microscope were imported to MATLAB based WorMachine^79^. A mask was generated for individual worms with default parameters from brightfield images. Worm length and GFP intensity for each mask were measured and compared using one-way ANOVA and Tukey HSD post hoc test in R packages.

##### Biosorter-based quantification

Animals on Day 3 adulthood were collected, washed, paralyzed, and surface bleached as described in the gut microbiome colonization steps, then transferred with 150 μl M9 buffer to a flat bottom 96 well plate (Costar 3370, Corning). Individual body size (time of flight, TOF) and level of GFP intensity were measured by a COPAS Biosorter (Union Biometrica) with a 250 micron flow cell and Sapphire488 laser at 310 volt and 1.0 pmt gain settings. Individual events were gated by a combination of TOF and extinction coefficient to filter adult animals from the population. GFP values normalized by TOF from each host strains and RNAi knockdown conditions were compared using one-way ANOVA and Tukey HSD post hoc test in R packages.

#### Developmental timing assays

Approximately 40 synchronized L1 worms were added to the plates containing BIGbiome mixtures or OP50. Animals were scored every 2 hrs for the number of adult animals on plate from 44 to 60 h at 20°C. Four replicates were scored for each condition, n > 100 animals were scored per strain/condition. Percentages of adults between the three microbiome types from the same time points were compared using one-way ANOVA and Tukey HSD post hoc test R packages.

### QUANTIFICATION AND STATISTICAL ANALYSIS

#### Random Forest analyses

To determine whether any of the microbial strains had predictive power over each of the microbiome types, we used random forest-based assessment in R using the randomForest package (https://www.rdocumentation.org/packages/randomForest/versions/4.6-14/topics/randomForest). Briefly, random forest is a supervised learning algorithm used to sort tested features into classes, and identify variables most strongly contributing to correct class prediction. A model was built with predictive variables of microbial relative abundance data and a response variable of microbiome type. Quality of the random forest was evaluated using the out-of-bag estimates of error rates (Mean Decrease in Accuracy) and confusion matrices. The data was split into training (60%) and testing sets (40%) and the model included 500 trees. A caveat of our analysis was that out-of-bag error (54.5%) and confusion matrix class errors (Type 1: 0.66, Type 2: 0.75, Type 3: 0.25) were high due to very uneven class sizes, but predictive taxa agreed with other assays nonetheless.

#### RNAseq analyses

See *Method* Details for explanation of software and programs used. In brief, reads were checked for quality with FASTQC, filtered for quality with bbmap, trimmed with bbduk, aligned with kallisto, and analyzed for differential expression with DESEQ2 ^77^. Genes were considered differentially expressed between microbiome types if the Benjamini-Hochberg adjusted p-value was less than 0.05. The work was completed locally using a Late 2013 Mac Pro (3.5 GHz 6-Core Intel Xeon E5) and software including Mac Terminal (fastqc, bbmap, bbduk, kallisto) and Rstudio (DESeq2).

#### Gene set enrichment analyses

The WormExp tool uses a statistical approach designed for gene list interpretation, EASE ^78^ to determine statistical significance. Default parameters were used and produced adjusted p-values based on False Discovery Rate (FDR) estimations. Adjusted P-values less than 0.05 were considered significant.

Rationale for statistical tools used within the WormCat tool are described in detail in Holdorf, *et al.,* ^29^. Briefly, WormCat produces Fisher’s exact test p-values. The method was chosen after providing few false positives without being too stringent in a randomized test of 100, 500, 1000, or 1500 genes. In our analyses, results were considered significant if the P-value score output from the WormCat online tool was less than 0.05.

#### Correlation analyses of microbial taxa abundance and gene expression

Pearson correlation was calculated between absolute abundance of each microbial taxa and expression of each gene for every strain of *C. elegans* using the ‘stats’ package within RStudio using default parameters. Correlations were considered significant if the Benjamini-Hochberg adjusted p-value was less than 0.05.

## SUPPLEMENTAL INFORMATION

### Supplemental Figures

**Figure S1.**
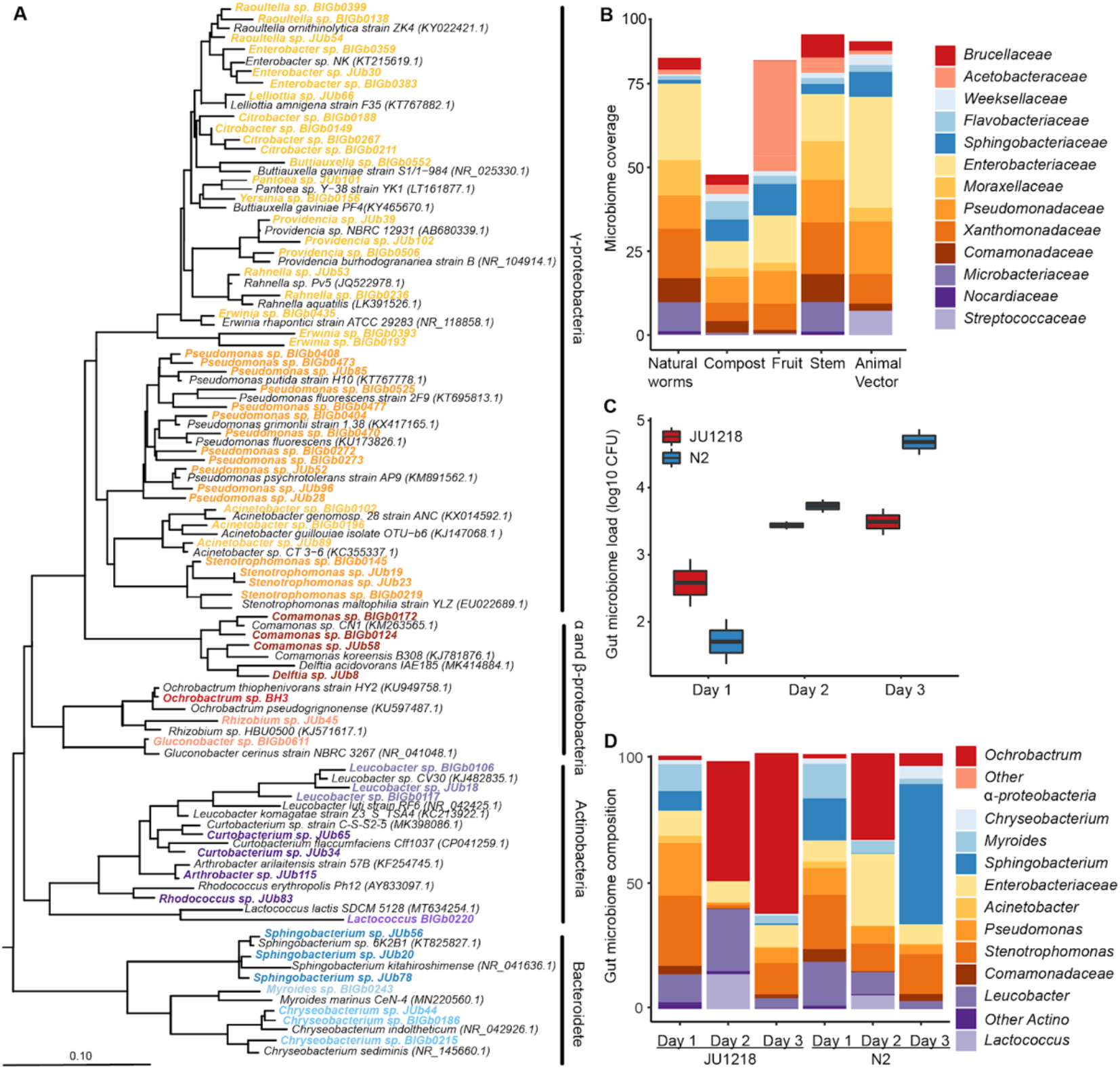
Establishment of a gut microbiome analysis pipeline in *C. elegans*. **A.** Phylogenetic tree is based on the full length of 16S rRNA genes of bacterial strains in this study using maximum likelihood method (Jukes–Cantor correction). BIGbiome strains are highlighted in bold and colored by corresponding genus. 16S rRNA gene sequence from *Synechococcus* sp. PCC7902 (AF216946) was used as an outgroup. **B.** BIGbiome strains represent from 49-90% of core microbiome families in *C. elegans* natural habitats [9]. **C.** Gut microbiome colonization increases from day 1 to day 3 adulthood in *C. elegans* N2 and JU1218 strains. **D.** Gut microbiome composition of N2 and JU1218 is similar on day 1 then diverges on Day 3 adulthood. Relative microbiome abundance was calculated from biological duplicates.

**Figure S2.**
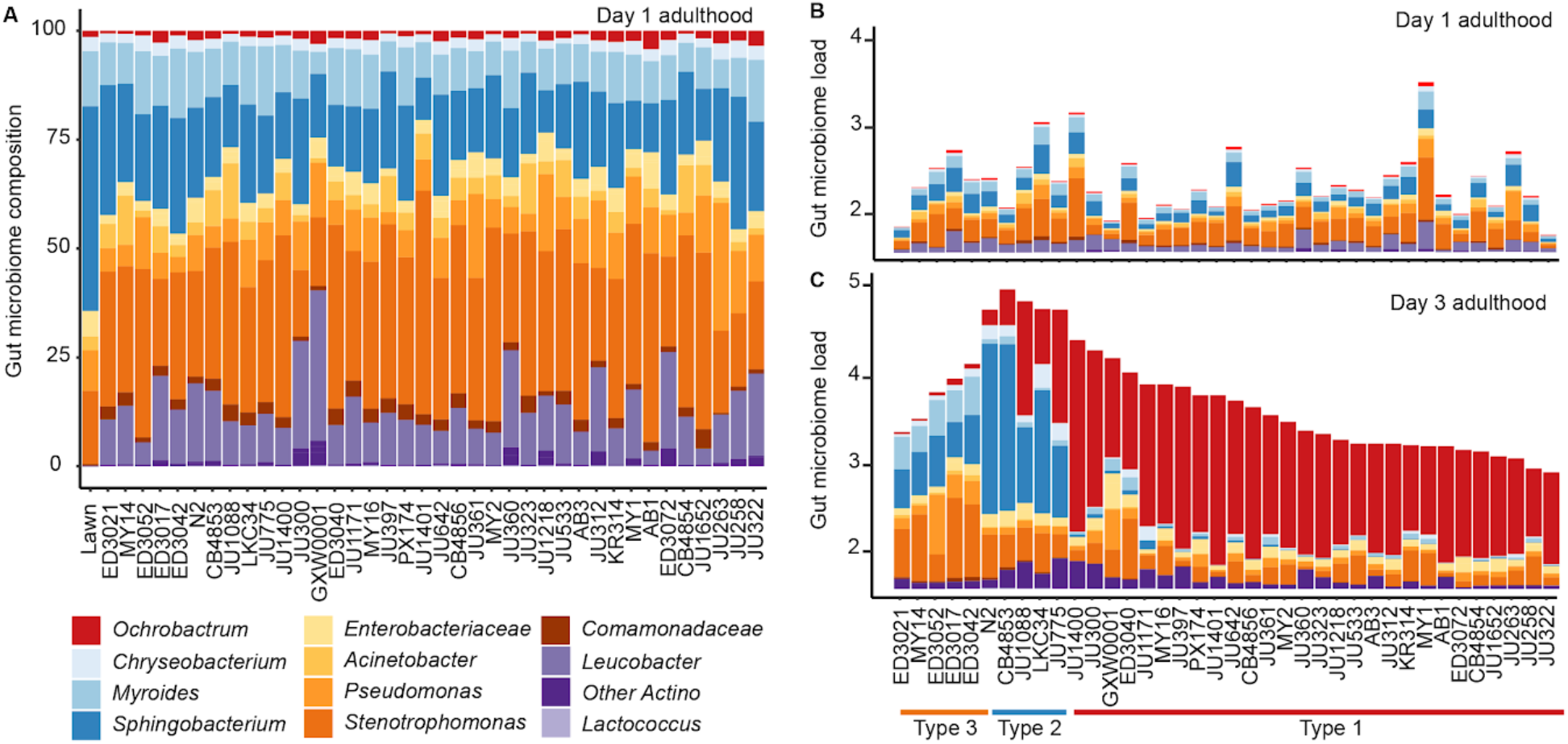
Minimal differences in *C. elegans* gut microbiomes at day 1 of adulthood. **A.** Gut microbiome composition of 38 *C. elegans* strains on Day 1 adulthood. Relative microbiome abundance was calculated from biological duplicates for each strain. **B.** Similar gut microbiome colonization levels of 38 *C. elegans* strains on Day 1 adulthood. Bar represented the mean calculated from biological duplicates for each strain. **C.** Gut microbiome colonization of 38 *C. elegans* strains on Day 3 adulthood. Type 2 strains, including lab-adapted N2 strain, carry on average a log higher gut microbiome load compared to Type 1 and 3 strains. Gut microbiome composition and load are calculated from the mean of biological duplicates for each strain.

**Figure S3.**
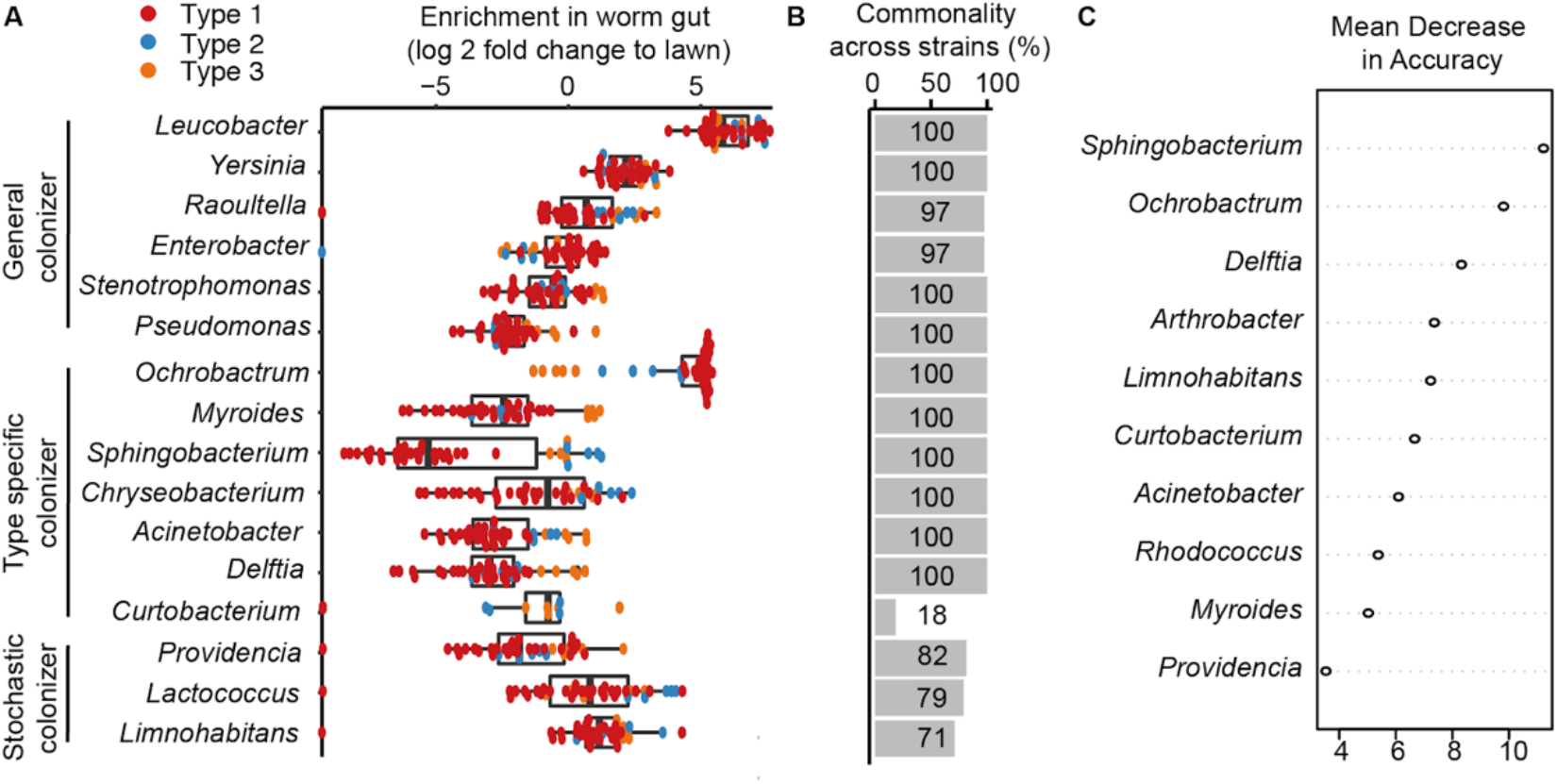
Microbial taxa that distinguish *C. elegans* gut microbiome types. **A.** Box-whisker plot of enrichment factors for microbial taxa for 38 *C. elegans* strains on day 3 adulthood colored by microbiome types. Enrichment factors for each microbial taxa were generated by log 2 transformation of fold changes of relative abundance in worm samples to BIGbiome lawn. **B.** Bar plot of commonality for each microbial taxa is calculated as the percentage of worm strains that was colonized by the corresponding microbial taxon at a minimum threshold of 0.01% in relative abundance. **C.** Variable importance plot (Random Forest) of 10 microbes with highest mean decrease in accuracy of microbiome type prediction upon their exclusion from a random forest model. Scores are determined using an out-of-bag error calculation.

**Figure S4.**
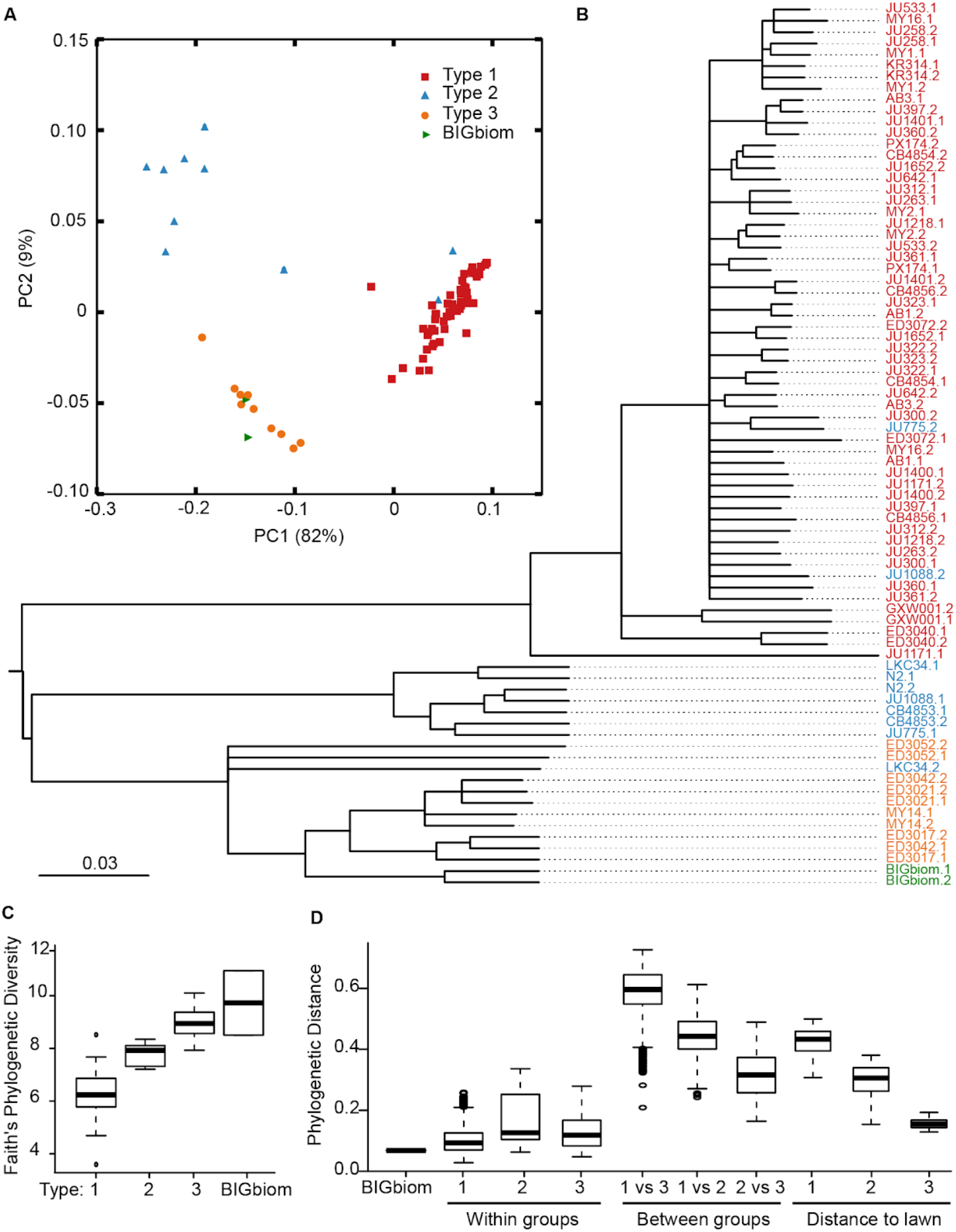
Microbiome diversity on day 3 of adulthood. **A.** PCoA of the microbiome composition of 38 *C. elegans* strains in Day 3 adulthood. **B.** Weighted jackknife clustering of microbiome composition of 38 *C. elegans* strains in Day 3 adulthood (colored by microbiome types). Replicates are shown under the same strain name. **C.** Box-whisker plot of Faith’s phylogenetic diversity for each of the three microbiome types and BIGbiome lawn. **D.** Box-whisker plot of weighted UniFRAC distance between the three microbiome types and BIGbiome lawn.

**Figure S5.**
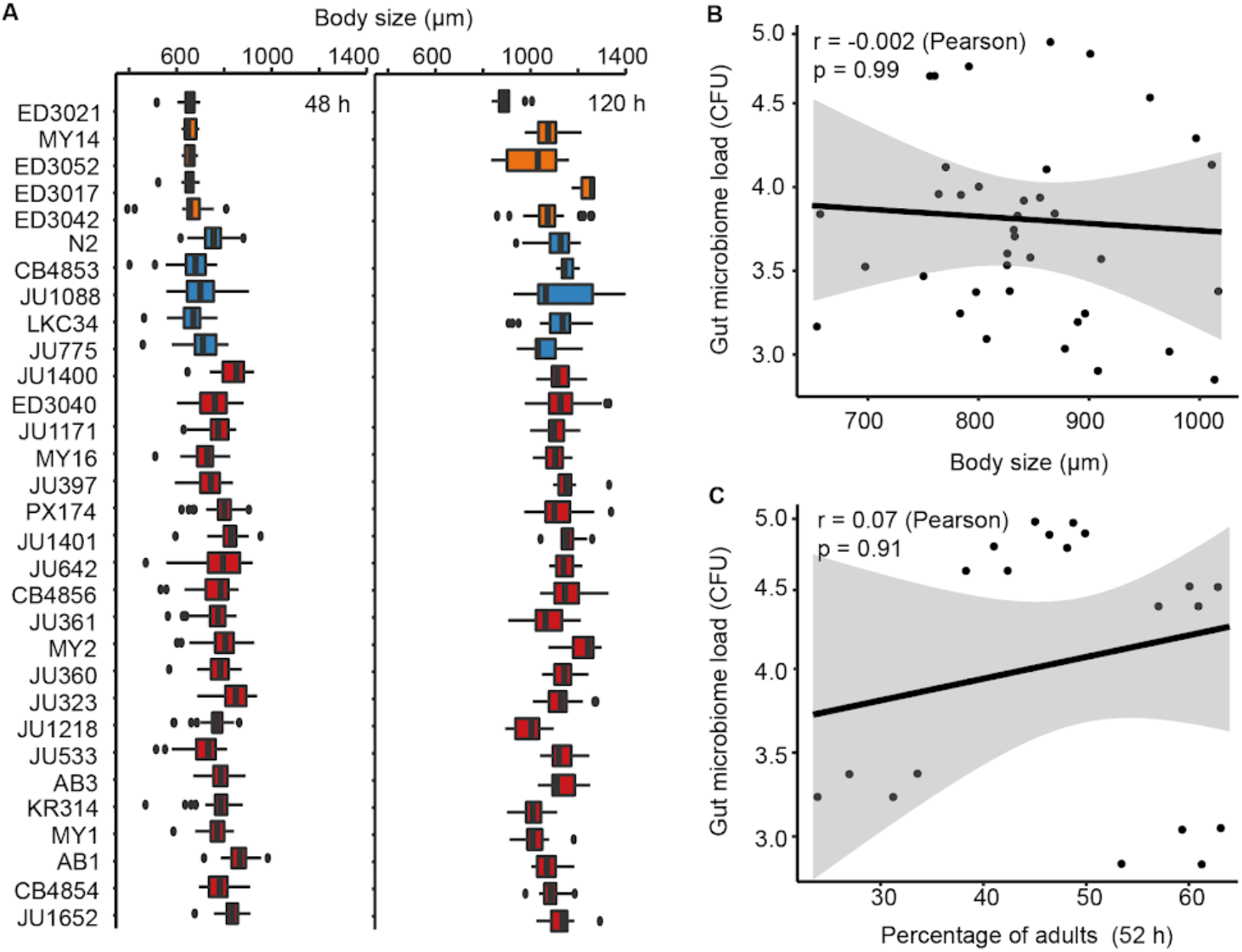
Gut microbiome load does not correlate with host development and body size. **A.** Box-whisker plot of body size of 38 *C. elegans* strains on day 1 and day 3 adulthood on BIGbiome. **B.** No correlation observed between gut microbiome colonization and percentage of adults at 52 h post L1s (r = −0.002, p = 0.99). **C.** No correlation observed between gut microbiome colonization and worm body size at 48 h post L1s (r = 0.07, p = 0.91).

**Figure S6.**
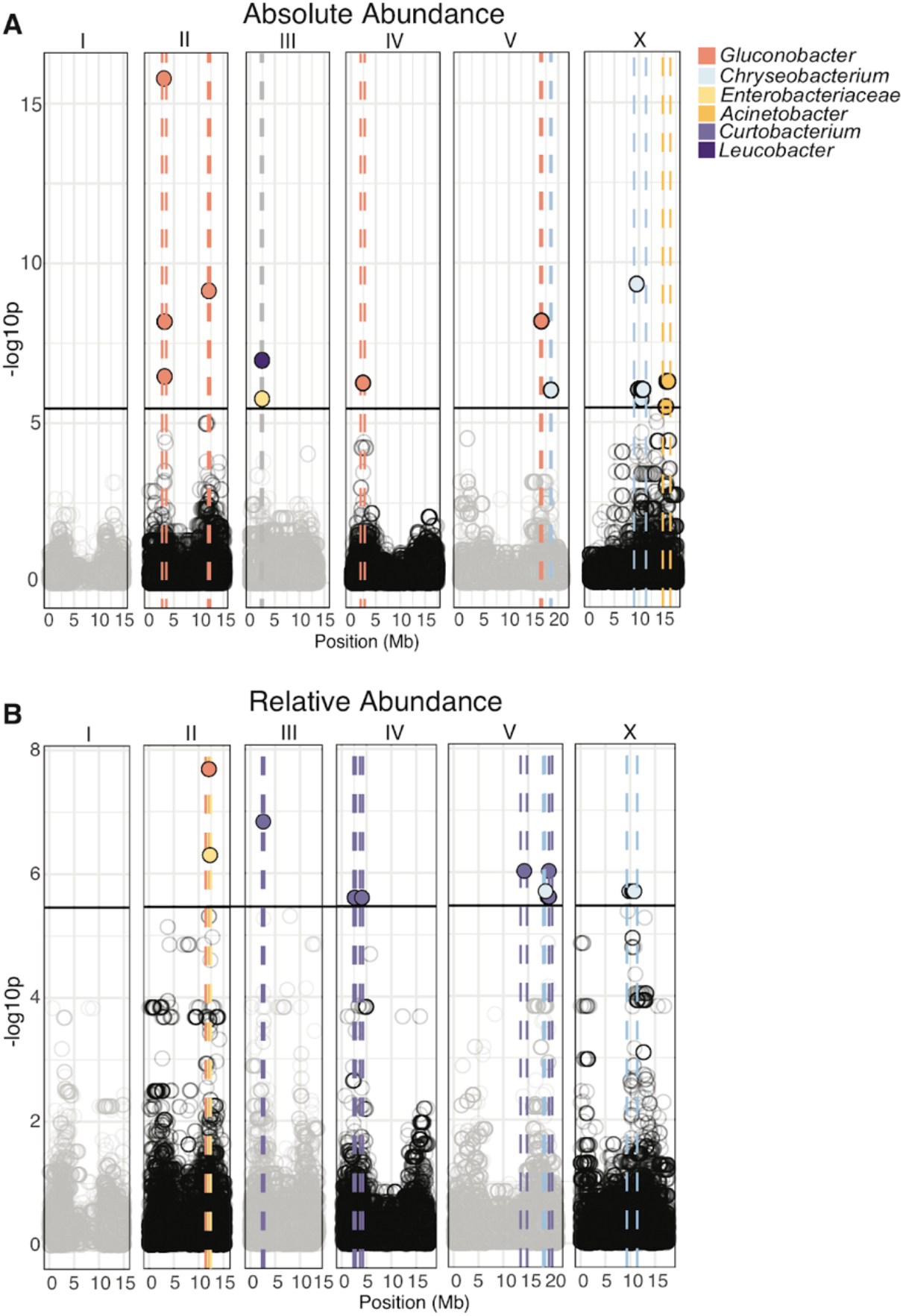
GWAS analyses identify genetic loci that are associated with gut microbiome abundance. **A.** GWAS plot for traits of absolute abundance of specific microbiome members. Points represent significance and genome region and are colored by microbe. Dashed lines indicate genomic region enriched for microbe-specific trait and are similarly colored by microbe. **B.** GWAS plot for traits of relative abundance of specific microbiome members. Points represent significance and genome region and are colored by microbe. Dashed lines indicate genomic region enriched for microbe-specific trait and are similarly colored by microbe.

**Figure S7.**
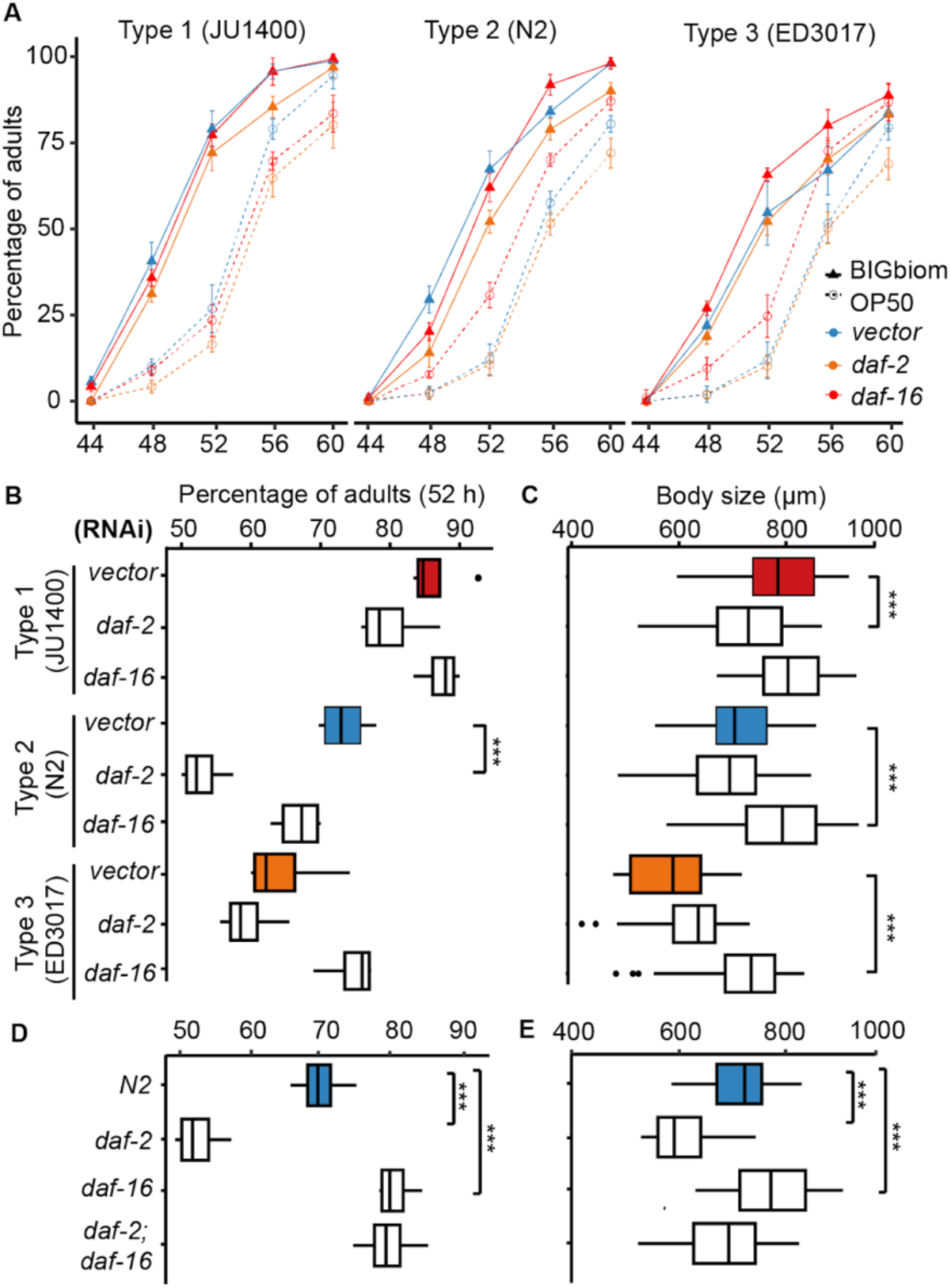
Development and body size of *daf-2* and *daf-16* mutants on BIGbiome. **A.** Developmental curves of *daf-2* and *daf-16* RNAi knockdown in representative microbiome type strains grown on the BIGbiome and *E. coli* OP50. Percentage of adults are represented as mean ± SD with 4 replicates for each condition. **B.** Box-whisker plot of adult percentages of vector(n=4), *daf-2*(n=4), *daf-16*(n=4) knockdown in representative strains of the three microbiome types at 52 h post L1 stage. **C.** Box-whisker plot of body size of vector, *daf-2*, *daf-16* knockdown mutants in representative strains of the three microbiome types at 48 h post L1 stage. **D.** Box-whisker plot of adult percentages of N2, *daf-2(e1370)*, *daf-16(mgDf50)*, and *daf-2(e1370);daf-16(mgDf50)* at 52 h post L1 stage (n=4 for each condition). **E.** Box-whisker plot of body size of N2, *daf-2(e1370)*, *daf-16(mgDf50)*, and *daf-2(e1370);daf-16(mgDf50)* at 48 h post L1 stage.

**Figure S8.**
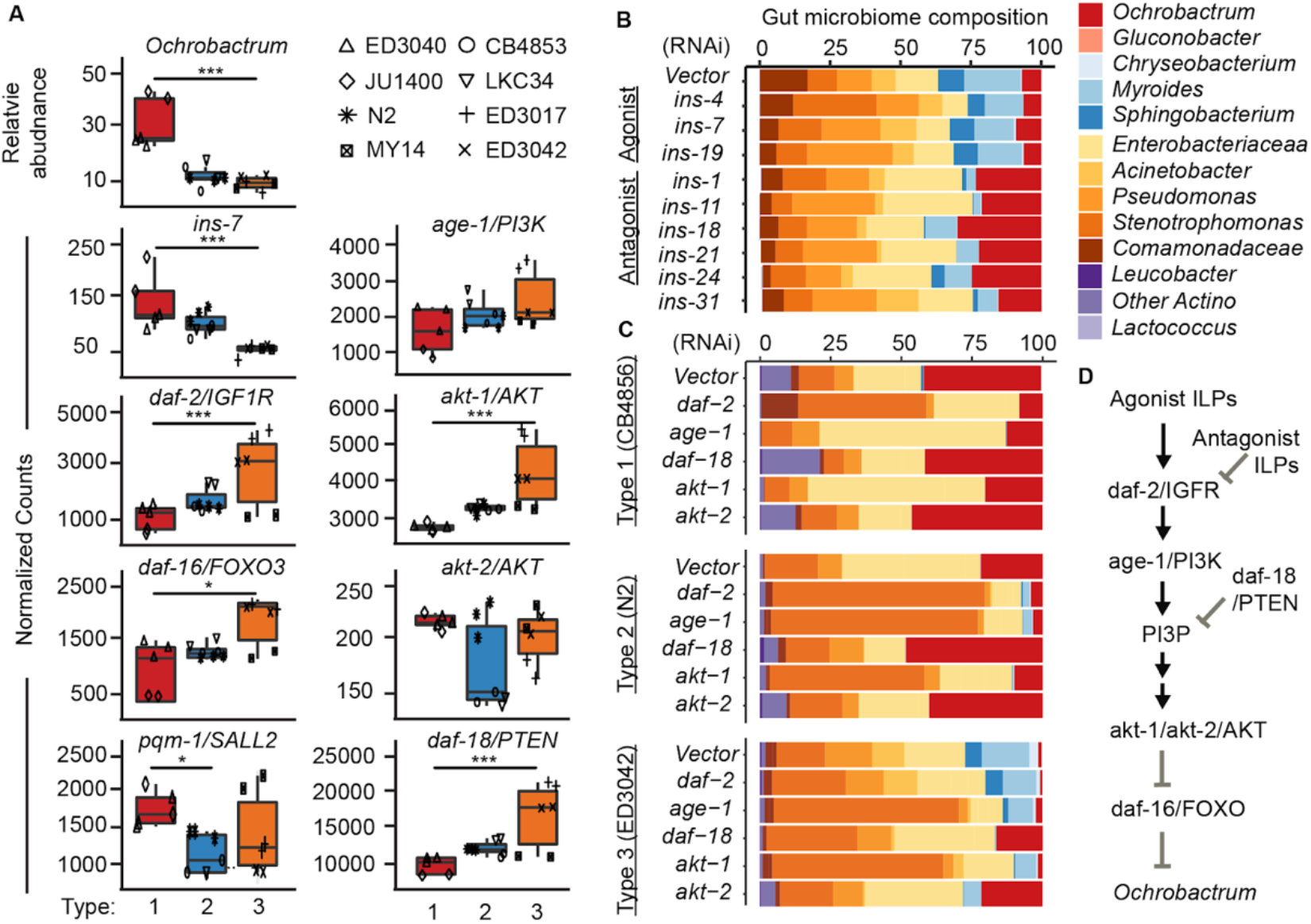
Key insulin signaling pathway genes that mediate recruitment of *Ochrobactrum*. **A.** Boxplots showing absolute abundance of *Ochrobactrum*, and expression of *ins-7, daf-2*, *daf-16, pqm-1, age-1, akt-1, akt-2,* and *daf-18* in each of the three microbiome types. **B.** Gut microbiome sequence showed *Ochrobactrum* colonization increased in the intestinal specific RNAi strain JM45 with *ins-11*(RNAi) and *ins-18*(RNAi). **C.** *Ochrobactrum* colonization in CB4856 (Type 1) decreased with *daf-2*(RNAi), *age-1*(RNAi), and *akt-1*(RNAi). In contrast, *Ochrobactrum* colonization increased in N2 (Type 2) and ED3042 (Type 3) with *daf-18*(RNAi) and *akt-2*(RNAi) shown by bulk gut microbiome sequence of the corresponding population. (**B,C**) Relative microbiome abundance is presented here as the mean of biological duplicates for each strain. **D.** Schematic diagram of insulin signaling pathway in *C. elegans*.

**Figure S9.**
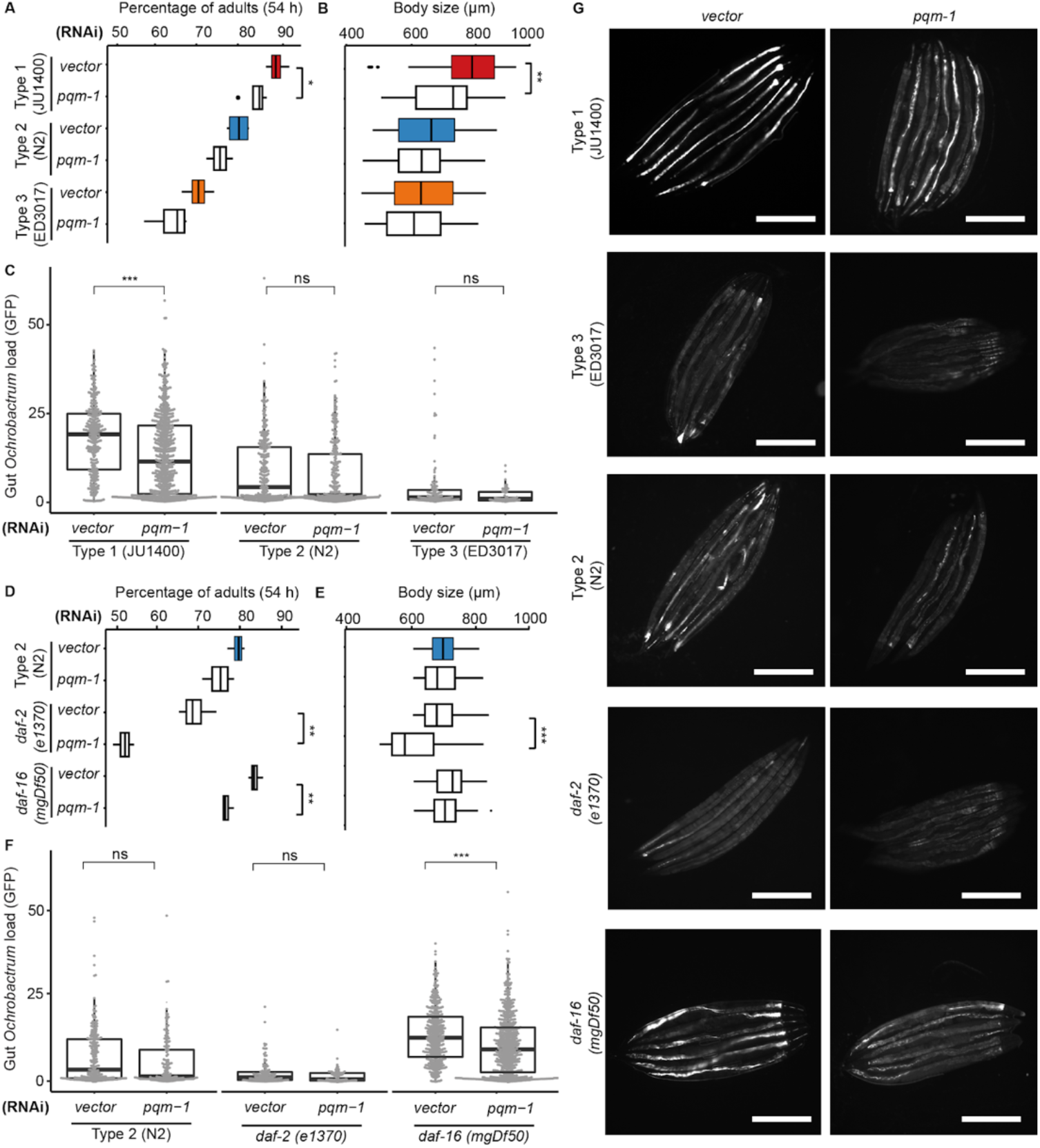
Further analyses of *pqm-1* on host development, body size, and *Ochrobactrum* colonization. **A.** Box-whisker plot of adult percentages of vector and *pqm-1* RNAi knockdown mutants in representative strains of the three microbiome types at 54 h post L1 stage. **B.** Box-whisker plot of body size of vector and *pqm-1* RNAi knockdown mutants in representative strains of the three microbiome types at 48 h post L1 stage. JU1400(vector): n=66, JU1400(*pqm-1*): n=85, N2(vector): n=79, N2(*pqm-1*): n=84, ED3017(vector): n=107, ED3017(*pqm-1*): n=68. **C.** Box-whisker plot of GFP-*Ochrobactrum* colonization of vector and *pqm-1* RNAi knockdown mutants in representative strains of the three microbiome types at 120 h post L1 stage. JU1400(vector): n=306, JU1400(*pqm-1*): n=657, N2(vector): n=290, N2(*pqm-1*): n=283, ED3017(vector): n=114, ED3017(*pqm-1*): n=57. **D.** Box-whisker plot of adult percentages of vector and *pqm-1* RNAi knockdown mutants in N2, *daf-2(e1370)*, and *daf-16(mgDf50)* at 54 h post L1 stage. **E.** Box-whisker plot of body size of vector and *pqm-1* RNAi knockdown mutants in N2, *daf-2(e1370)*, and *daf-16(mgDf50)* at 48 h post L1 stage. N2(vector): n=206, N2(*pqm-1*): n=267, *daf-2*(vector): n=300, *daf-2*(*pqm-1*): n=109, *daf-16*(vector): n=136, *daf-16*(*pqm-1*): n=179. **F.** Box-whisker plot of GFP-*Ochrobactrum* colonization of vector and *pqm-1* RNAi knockdown mutants in N2, *daf-2(e1370)*, and *daf-16(mgDf50)* at 120 h post L1 stage. N2(vector): n=237, N2(*pqm-1*): n=164, *daf-2*(vector): n=122, *daf-2*(*pqm-1*): n=75, *daf-16*(vector): n=412, *daf-16*(*pqm-1*): n=621. (**B**,**E**) n represents the number of individual animals quantified by microscopic images.(**C**,**F**) n represents the number of individual animals quantified by Biosorter. P-values were generated from student’s t-test (***p<0.001, **p<0.01,*p<0.05). **G.** Representative images of GFP-*Ochrobactrum* colonization of vector and *pqm-1* RNAi knockdown mutants in Type 1(JU1400), Type 3(ED3017), Type 2(N2), *daf-2(e1370)*, and *daf-16(mgDf50)* at 120 h post L1 stage (Bar = 500 μm).

### Supplemental Tables and Files

**Table S1. List of bacterial strains used in this study.** This table contains metadata related to bacterial strains in BIGbiome001, including isolation origin, taxonomy, and 16S rRNA sequences.

**Table S2. List of *C. elegans* strains used in this study.** This table contains metadata related to *C. elegans* strains grown on BIGbiome001, including isolation locations and substrates.

**Table S3. Enrichment and depletion of microbial taxa within the *C. elegans* gut compared to lawns.** This table lists the average fold changes of gut microbial abundance normalized to their abundance in the BIGbiome lawn on day 3 adults of 38 wild *C. elegans*, highlighting host effects in enriching or depleting microbial colonization in the gut environment. Commonality for each taxa is calculated based on the number of minimum presence (>0.01%) across 38 wild *C. elegans. C. elegans* strains in columns are grouped by gut types with the same color scheme as the main figures.

**Table S4. Pearson correlation of gut microbial abundance with host body size and developmental growth rate.** This table lists the Pearson correlation coefficient between microbial abundance on day 3 adults and two host phenotypes (developmental growth rate/body size). p-value was computed at the level of 95% confidence interval. *Ochrobactrum* (highlighted in red) is the only microbial taxa with positive correlation with both host phenotypes (p<0.05). 9 other microbial taxa (highlighted in blue) show negative correlation with both host phenotypes (p<0.05).

**Table S5. Summary of GWAS loci associated with bacterial taxa.** This table contains direct output information from the CeNDR GWAS tool including marker, chromosome, position, user-provided trait (microbe and abundance, in this case), log10(p-value), significance threshold (BF), binary signifier if the trait was above the BF score, *C. elegans* strain, value, allele, var.exp, start and stop position, peak position, peak identification, and interval size (see reference 82 for more information about the CeNDR tool and information outputs). Tabs are separated by microbe and type of abundance (relative or absolute).

**Table S6. Genes differentially expressed between microbiome types.** This table contains output information from the DESeq2 results outputs for each pairwise comparison of microbiome types. Each row corresponds to a gene, with the pair of gut types being compared, and information on the base mean expression level, log2(fold change), and uncorrected and corrected p-values (see reference 88 for more information about the DESeq2 tool and information output files).

**Table S7. Raw data associated with figures and tables.** This table contains raw data to support the figures and tables in the manuscript. A summary tab describes data types and corresponding figure panels.

**File 1. Code and scripts used in the analysis of datasets.** The provided archive file contains code and scripts for all analyses in this study, including those for overall microbiome compositional analyses (‘Microbiome_analysis_scripts.txt’), predictive taxa for microbiome types (‘RandomForest_Script.txt’), processing and analysis of RNAseq datasets (‘Kallisto_bbmap_bbduk_Script.txt’ and ‘DifferentialExpression_Script.txt’), correlation of microbial taxa with gene expression profiles (‘MicrobialAbundance_vs_GeneExpression_Correlation_Script.txt’) and figure generation in R environment (‘R_plot_figures.txt).

