## Supplemental File 1 for "Natural genetic variation drives microbiome selection in the *Caenorhabditis elegans* gut": MicrobialAbundance_vs_GeneExpression_Correlation_Script.txt

###################################################################################

#Correlation Analysis and p-value calculation (RStudio)

#Functions


#Function to calculate Pearson correlation scores between gene expression and absolute abundance measurements of a designated microbe
correlation<-function(expr, abu, microbe, outFile) {    outFile[[microbe]]<-NA #User inputs include file containing expression data, file containing absolute abundance scores, variable that should be the same as the column name of the microbe of interest in the absolute abundance scores file, and the name of the output file

correlation<-function(expr, abu, microbe, outFile) {    outFile[[microbe]]<-NA
    for(i in 1:18715) {

        rownames(outFile)[[i]]<-colnames(expr)[[i]] #Name rows of output file based on column names of the expression data file (should correspond to gene name)

        pearsonScore<-cor(abu[,microbe], expr[,i]) #Calculate Pearson correlation score 

        outFile[i,microbe] <- pearsonScore #Print Pearson correlation score to output file

    }    

    return(outFile)}

#Function to calculate Benjamini-Hochberg corrected p-value of the Pearson correlation between between gene expression and absolute abundance measurements of a designated microbe

PearsonCorrelation_pvals<-function(expr, abu, microbe, outFile) {    outFile[[microbe]]<-NA #User inputs include file containing expression data, file containing absolute abundance scores, variable that should be the same as the column name of the microbe of interest in the absolute abundance scores file, and the name of the output file

    for(i in 1:18715) {

         rownames(outFile)[[i]]<-colnames(expr)[[i]] #Name rows of output file based on column names of the expression data file (should correspond to gene name)

         pval<-cor.test(x=as.matrix(abu[,microbe]), y=as.matrix(expr[,i]), adjust = "BH") #Calculate Benjamini-Hochberg adjusted p-value

         pearsonScore_pval<-pval$p.value #Put the p-value into a variable

         outFile[i,microbe] <- pearsonScore_pval #Print Benjamini-Hochberg adjusted p-value to output file

     }    

     return(outFile)

 }


RNAseq_correlations<-data.frame(matrix(nrow = 18715))  #Initialize the data frame 

RNAseq_correlations<-correlation(enterotype_Adrien_DEseq2_ddsTxi_counts_t, RNAseq_AbsAbundance, [microbe], RNAseq_correlations) #Run correlation function

RNAseq_correlations_pvals<-PearsonCorrelation_pvals(enterotype_Adrien_DEseq2_ddsTxi_counts_t, RNAseq_AbsAbundance, [microbe], RNAseq_correlations_pvals) #Run function to calculate BH-adjusted p-values
