## Supplemental File 1 for "Natural genetic variation drives microbiome selection in the *Caenorhabditis elegans* gut": Microbiome_analysis_scripts.txt

#1. Check map files, if necessary
validate_mapping_file.py -m cemicro_map.txt -o validate_mapping_file_output/cemicro_corrected.txt

#2. Split barcoded libraries into one file and quality filter
split_libraries_fastq.py -i R1_001.fastq.gz,R2_001.fastq.gz -b I1_001.fastq.gz -m cemicro_corrected.txt --rev_comp_mapping_barcodes -q30 --retain_unassigned_reads -o seqs.fna 
	
#3. Remove chimeras with Usearch61
identify_chimeric_seqs.py -m usearch61 -i seqs.fna -r gg_13_8_otus/rep_set/99_otus.fasta --percent_id_usearch61 0.99 -o usearch61_chimera_checking/chimeras.txt

filter_fasta.py -f seqs.fna -o seqs_check.fna -s /usearch61_chimera_checking_5/chimeras.txt -n

#4. Generate ASV from deblur
source activate deblurenv
deblur workflow --seqs-fp seqs_check.fna --output-dir output -t 250 --neg-ref-fp --pos-ref-fp BIGbiome.fasta

##the output folder generates a file reference-hit.biom, which contains the output of positive filtering of the sOTUs
##the output folder also generates a file reference-hit.seqs.fa containing the sequence information of the sOTUs. This file is used to build a maximum likely hood tree named rep_set.tre in Mega, which is used in beta diversity analysis below.  

#6. Run diversity analyses 
core_diversity_analyses.py -i reference-hit.biom -o $PWD/core_output -m cemicro_corrected.txt -c Strains -t rep_set.tre -e 3000

jackknifed_beta_diversity.py -i reference-hit.biom  -o bdiv_jk100 -e 3000 -m cemicro_corrected.txt -t rep_set.tre
