## Supplemental File 1 for "Natural genetic variation drives microbiome selection in the *Caenorhabditis elegans* gut": RandomForest_Script.txt

###################################################################################

#Random Forest (RStudio) [see followed tutorial: https://web.archive.org/web/20191211192257/http://dni-institute.in/blogs/random-forest-using-r-step-by-step-tutorial/]

#Examine properties of dataset - percent of samples falling in each enterotype

table(metadata_trimmed$enterotype)/nrow(metadata_trimmed)


#Split data into training and validation sets - see that both training and validation sets have similar distributions into the three enterotypes to one another and the original data 

RF_sample.ind<-sample(2, nrow(metadata_trimmed), replace = T, prob = c(0.6,0.4))

RF_cross.sell.dev<-metadata_trimmed[RF_sample.ind==1,]

RF_cross.sell.val<-metadata_trimmed[RF_sample.ind==2,]

table(RF_cross.sell.val$enterotype)/nrow(RF_cross.sell.val)

table(RF_cross.sell.dev$enterotype)/nrow(RF_cross.sell.dev)


#Change the data type of enterotype to factor

metadata_trimmed$enterotype<-as.factor(metadata_trimmed$enterotype)


#Make the formula

RF_varNames<-names(RF_cross.sell.dev)  

RF_varNames<-RF_varNames[!RF_varNames %in% c("enterotype")]         #exclude the response variable

RF_varNames1<-paste(RF_varNames, collapse = "+")                    #add "plus" sign between predictive variables 


#Add response variable and convert to formula object

RF_rf.form<-as.formula(paste("enterotype", RF_varNames1, sep = " ~ "))


#Find optimal number of variables to split at each node

RF_enterotype.bestmtry<-tuneRF(metadata_trimmed[-1], metadata_trimmed$enterotype, ntreeTry=500, stepFactor = 1.5, improve = 0.01, trace = TRUE, plot = TRUE, doBest = FALSE)


#Build 500 decision trees and evaluate 

RF_cross.sell.rf <- randomForest(RF_rf.form, RF_cross.sell.dev, ntree=500, importance=T)

print(RF_cross.sell.rf)


#Predict response variable using random forest

RF_enterotype.dev$predicted.response<-predict(RF_enterotype.rf, RF_enterotype.dev)


#Plotting scatterplot matrix of most discriminative features 

pairs(RNAseq_AbsAbundance[,c(8, 11, 23, 9)], col=colors, pch=19)
